# Genetically controlled mtDNA editing prevents ROS damage by arresting oxidative phosphorylation

**DOI:** 10.1101/2020.11.20.391110

**Authors:** Simon Stenberg, Jing Li, Arne B. Gjuvsland, Karl Persson, Erik Demitz-Helin, Carles Gonzalez Peña, Jia-Xing Yue, Ciaran Gilchrist, Timmy Ärengård, Payam Ghiaci, Lisa Larsson-Berglund, Martin Zackrisson, Johanna L. Höög, Mikael Molin, Gianni Liti, Stig W. Omholt, Jonas Warringer

## Abstract

Deletion of mitochondrial DNA in eukaryotes is mainly attributed to rare accidental events associated with mitochondrial replication or repair of double-strand breaks. We report the discovery that yeast cells arrest harmful intramitochondrial superoxide production by shutting down respiration through genetically controlled deletion of mitochondrial oxidative phosphorylation genes. We show that the regulatory circuitry underlying this editing critically involves the antioxidant enzyme superoxide dismutase 2 and two-way mitochondrial-nuclear communication. While mitochondrial DNA homeostasis is rapidly restored after cessation of a short-term superoxide stress, long-term stress causes maladaptive persistence of the deletion process, leading to complete annihilation of the cellular pool of intact mitochondrial genomes and irrevocable loss of respiratory ability. Our results may therefore be of etiological as well as therapeutic importance with regard to age-related mitochondrial impairment and disease.

**One-Sentence Summary:** Genetically controlled editing of mitochondrial DNA is an integral part of the yeast’s defenses against oxidative damage.

## Main Text

Mitochondrial impairment is strongly associated with aging (*1*) and the pathogenesis of age-related human diseases, including Alzheimer’s disease (*2*), Parkinson’s disease (*3*), the deterioration of skeletal and cardiac muscle (*4*), and macular degeneration (*5*). Mitochondrial DNA (mtDNA) deletions are perceived to contribute markedly to this impairment (*6*). The general conception is that mtDNA deletions are caused by accidental events associated with replication or repair of double-strand breaks (*7*). However, considering that mtDNA deletions are prone to cripple oxidative phosphorylation (*8*) (OXPHOS) it is conceivable that mtDNA deletion is also under genetic control, the main rationale being that compared to a mitophagic response (*9*–*13*), it would serve as a less costly defense mechanism against an abrupt increase in intramitochondrial superoxide production inundating the primary antioxidant defenses (*14*–*16*). The disclosure of an additional genetically controlled defense layer against superoxide damage, situated between the primary antioxidant defenses and mitophagy, would bring a fresh perspective to what causes mitochondrial impairment and how it can be mitigated. This motivated us to search for the existence of such a regulatory layer in budding yeast (*Saccharomyces cerevisiae*).

### Swift adaptation to enhanced superoxide production

To avoid possible confounding effects of domestication (*17*), we exposed wild haploid yeast cell populations expanding clonally on glucose to the mitochondrial superoxide anion (O_2_^•−^) generator and redox cycler paraquat (N,N-dimethyl-4-4′-bipiridinium dichloride) (*18*). As OXPHOS complex III and mitochondrial NADPH dehydrogenases donate electrons to paraquat, which passes these on to O_2_ (*18, 19*), this mode of O_2_^•−^ generation is a good proxy for the *in vivo* situation (*20*). By titration of the paraquat dose, we ensured that the O_2_^•−^ production was beyond the regulatory reach of the primary antioxidant defenses, while not severely compromising cellular function (fig. S1, supplementary text 1).

We then observed how 96 clonally reproducing yeast cell populations on solid agar medium adapted to the chosen paraquat dose in terms of change in cell doubling time over approximately 240 generations (corresponding to 50 growth cycles). To provide a comparative data set, we similarly exposed a total of 672 populations to seven other stressors not explicitly challenging mitochondrial function (table S1). All 96 populations exposed to paraquat adapted much faster than every other population exposed to the other stressors (Fig. 1A). Within the first 10 generations, cell populations reduced their doubling time by on average 106 min, corresponding to 49.3% of the maximum possible reduction. Thenceforth, adaptation entered a second phase where the reduction in cell doubling time progressed much slower until it plateaued after 75 generations at 72.6% of the maximum possible reduction (mean) (Fig. 1A).

**Figure 1.**
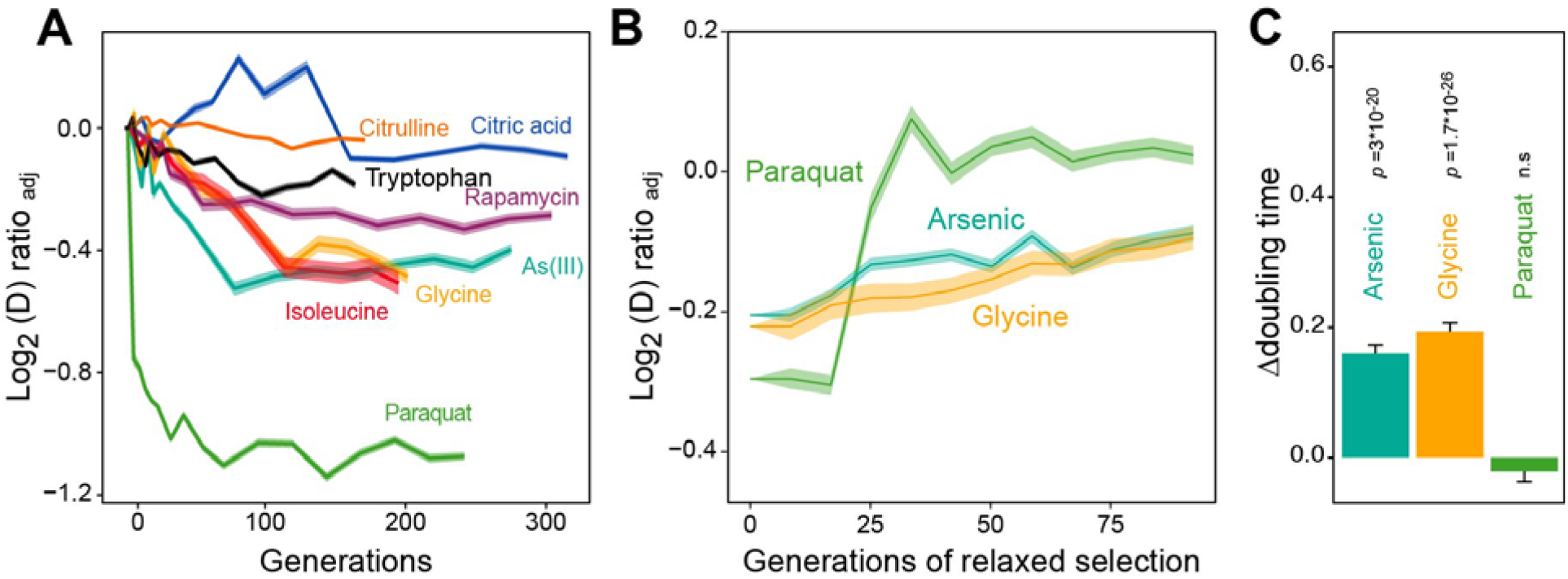
Distinct adaptation to paraquat. (**A)** Mean temporal adaptive response to paraquat and seven other stressors. *y*-axis shows log_2_ fold reduction in cell doubling time (h) from pre-stress, adjusting for plate, position and pre-culture effects. 96 populations for each stressor (*n=*6). Shade: S.E.M. (**B**) Loss of the acquired adaptation as a function of number of cell generations after release from the selection pressure. Colored lines: mean of 96 populations (each measured at *n=*5). Shade: S.E.M. The populations were released from stress after reaching 70-90% of their endpoint (*t*_*50*_) adaptation. (**C**) The difference in cell doubling time (h) in a no-stress environment between 96 populations (each measured at *n=*5) having achieved 70-90% of their endpoint adaptation to paraquat, arsenic and glycine, respectively, and the founder population. The difference reflects the selective advantage of losing the acquired adaptations when the populations are no longer exposed to stress. Error bars: S.E.M.

A numerical model of the adaptation process, combining population genetics and population dynamics concepts with experimental data on nuclear mutation rates and effect sizes of *de novo* point mutations and aneuploidies (*21*), was completely unable to reproduce the observed extraordinarily swift response to paraquat (fig. S2).

After being exposed to paraquat over a few growth cycles before being placed in a paraquat-free medium, all 96 cell populations retained their acquired tolerance to paraquat (mean reduction in cell doubling time: 106 min) for only 1-2 growth cycles before abruptly losing it (Fig. 1B). When employing the same experimental procedure to 96 cell populations from each of the two other environments to which adaptation was also fast (arsenic and glycine), we found that despite the presence of a much stronger Darwinian counterselection (Fig. 1C), these populations lost their acquired adaptations more slowly and gradually (Fig. 1B). While a classical Darwinian mutation/selection-based adaptive process could reasonably explain the data for seven of the eight tested stressors, we concluded that the data on paraquat could hardly be reconciled with such a process.

### Mitophagy is not responsible for the swift first adaptation phase

We next probed whether a mitophagic response was responsible for the swiftness of the first adaptation phase. As mitochondrial fragmentation is a well-documented prelude to canonical mitophagy (*22*), we assayed mitochondrial morphology before and after paraquat exposure by confocal and electron microscopy. We observed a rapid shift (<5 h) from a tubular to a fragmented mitochondrial organization (Figs. 2A, B, fig. S3A) as well as a rapid reversal (<5 h) to a tubular organization after removal of stress, consistent with the notion that mitochondrial O_2_^•−^ generation influences the mitochondrial fission and fusion dynamics (*23, 24*). DNA staining indicated that the mitochondrial fragments generally contained mtDNA after 5 h of paraquat exposure (fig. S3B). The mitochondrial volume remained near pre-stress levels with at the most a marginal reduction after 77-79 h of paraquat exposure (*22*) (Figs. 2A, B). Moreover, cell populations (*n=*16) lacking Atg32, a key component of canonical mitophagy (*25*), adapted to paraquat over ∼80 generations as wild type populations (Fig. 2C). These observations led us to conclude that the initial swift adaptation to paraquat was not likely to depend on canonical mitophagy.

**Figure 2.**
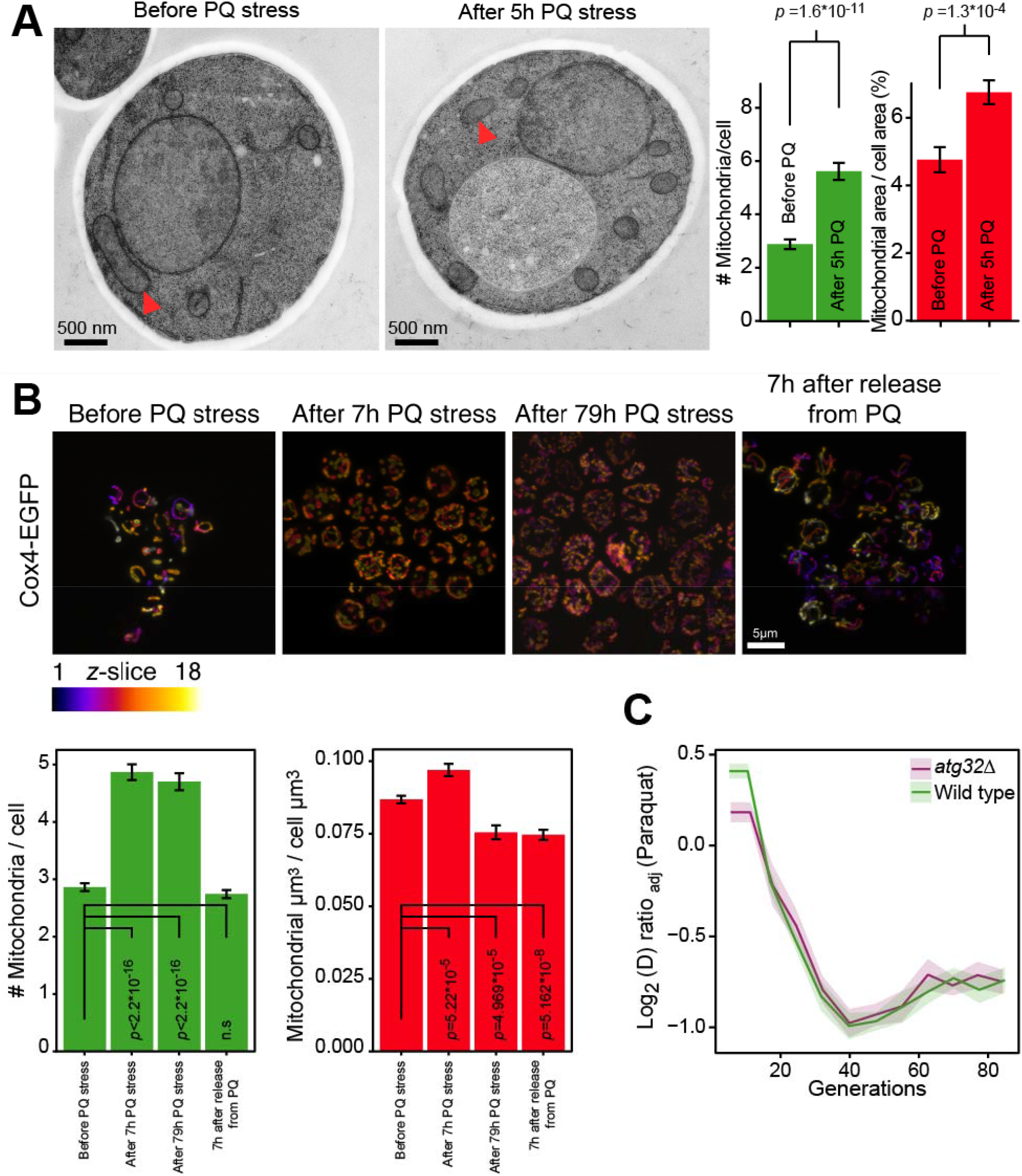
Mitochondrial fragmentation precedes the swift adaptation to paraquat (PQ). (**A**) EM microscopy of cells before (panel 1) and after short-term stress (panel 2, red arrowheads mark representative mitochondria). Panel 3 shows the number of mitochondria per imaged cell (left) and the imaged cell area occupied by mitochondrial area (%) (right), used as proxy for mitochondrial volume. Error bars: S.E.M. (*n=*100 cells). *p*-values: Welch two-sided t-test. (**B**) Confocal microscopy of cells with a Cox4-GFP mitochondrial tag. Color: *z*-dimension (yellow=front, purple=back; 18 slices). *Lower left diagram:* Number of mitochondria per imaged cell. *Lower right diagram:* Mean sum of mitochondrial volume as a fraction of cell volume. Error bars: S.E.M. (*n*=473-910 cells). *p*-values: Welch two-sided t-test (**C**) Adaptation of *atg32Δ* and wild type populations to paraquat. Shade: S.E.M. (*n*=16 populations, each measured at *n=*6).

### mtDNA loss drives the first adaptation phase

We then assayed the copy numbers of mtDNA genes in nine paraquat-adapting cell populations by qPCR. In line with short read sequencing results from five populations (Fig. 3A, supplementary text 2), we found that cells lost parts of their mtDNA during the early adaptation phase. In addition, the qPCR data showed that the lost segments were unevenly distributed across the mitochondrial genome: all nine cell populations lost one or more segments within the mtDNA region spanning *COX1* to *VAR1*, while a few also lost the 21S rRNA and *COX2* rapidly thereafter (Fig. 3B, fig. S4). Cells retained the mtDNA segments that were not lost at near founder levels throughout the early adaptation phase, which in all nine cases encompassed *COX3-RPM1* and 15S rRNA. Since the mtDNA coverage prior to paraquat adaptation was perfectly even (fig. S5A), the observed mtDNA loss was clearly induced by paraquat.

**Figure 3.**
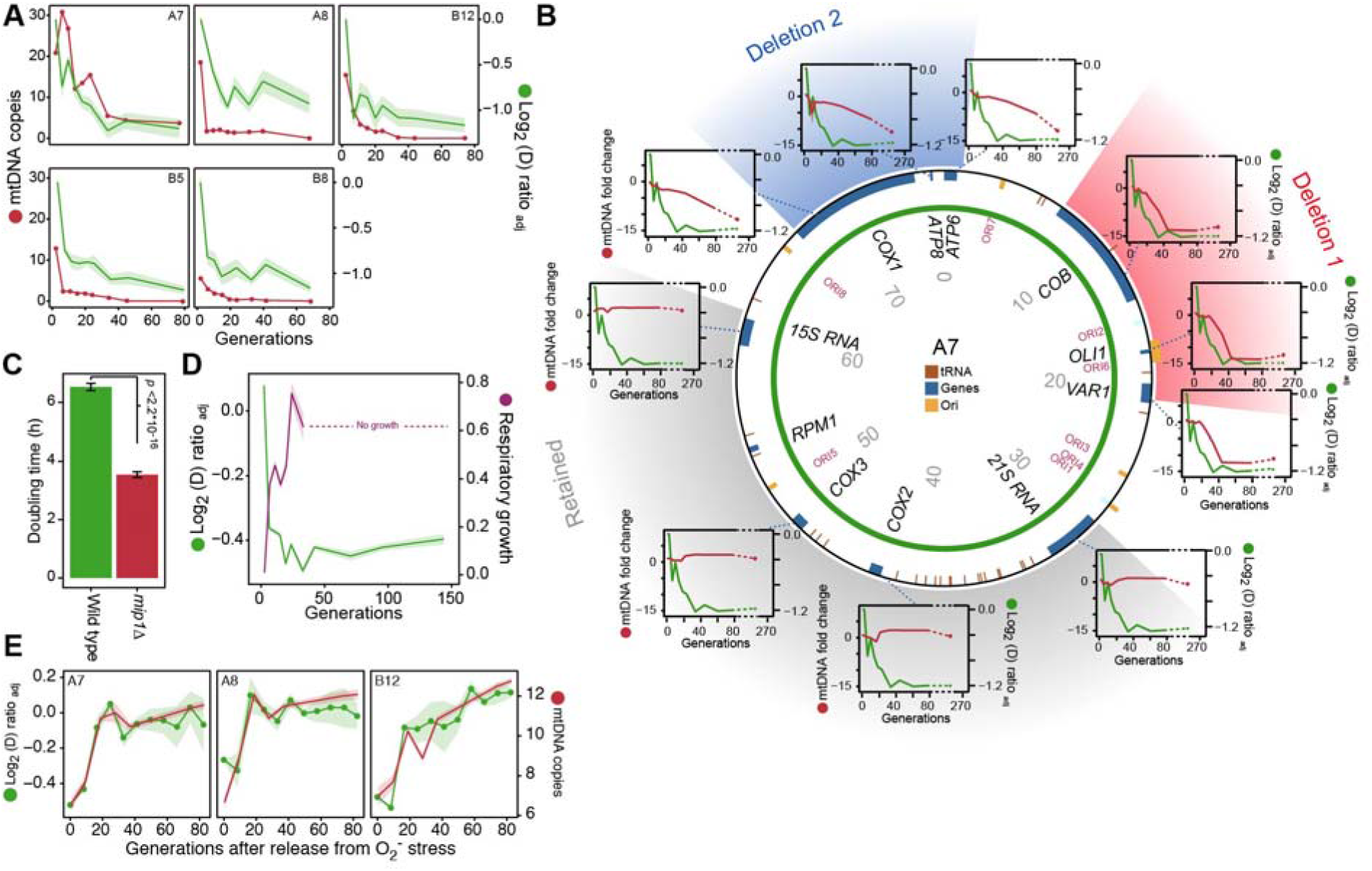
mtDNA editing causes the early adaptation to paraquat. (**A**) mtDNA copy number change (left *y*-axis, red line, median coverage relative to the haploid nuclear genome) during paraquat adaptation (right y-axis, green line, *n=*6) for 5 populations (panels). Shade: S.E.M. (**B**) mtDNA deletions associate with the paraquat adaptation. *Circle:* mtDNA (77 kb) before exposure to paraquat. Genes, origins of replication and position (kb) are indicated. *Coloured fields:* mtDNA deletions with concerted copy number change. *Diagrams:* mtDNA copy number change (left *y*-axis, purple line) of individual mtDNA genes during adaptation (right *y*-axis, green line) in population A7. Shade: S.E.M. (*n=*3). (**C**) Doubling time (*h*) of wild type and *mip1Δ* cell populations in paraquat. Error bars: S.E.M. (*n=*191). *p*-values: Welch two-sided t-test. (**D**) The early-phase paraquat adaptation coincides with the loss of respiratory (glycerol) growth. Shade: S.E.M. 96 populations, each measured at *n=*5. Broken line indicates no growth (cell doubling time>24h). (**E**) Recovery of the copy number of deleted mtDNA (right *y*-axis, red line) after release from 6 generations of paraquat exposure coincides with loss of the early-phase paraquat adaptation (left *y*-axis, green line, shade = S.E.M (*n*=15)) in populations A7, A8 and B12.

Anticipating that the observed mtDNA loss caused a reduction in OXPHOS activity, and thus in mitochondrial O_2_^•−^ production, we exposed *ρ*^0^ cell populations (*n=*12) (supplementary text 3), obtained by deletion of the sole mitochondrial DNA polymerase Mip1 (Pol γ homolog) (*26*), to the same concentration of paraquat as was used in the initial adaptation experiments. These *mip1Δ* cells were highly resistant to paraquat compared to wild type cells (Fig. 3C). Moreover, exposing the pre-adapted *mip1Δ* cell populations to paraquat for 10 growth cycles caused no or marginal additional improvement in growth (fig. S5B). To substantiate these findings, we cultivated cell populations exposed to short-term paraquat stress on the respiratory growth medium glycerol. The early adaptation correlated almost perfectly with loss of the capacity for respiratory growth (Fig. 3D), and the rapid loss of the acquired adaptations after removal of stress coincided with the restoration of the ability for respiratory growth (fig. S5C). We next repeated the stress release experiment on the five sequenced populations, and tracked by short-read sequencing the temporal change in copy numbers of the lost mtDNA segments. As expected, the five populations quickly restored their ability for respiratory growth after removal of paraquat, and this coincided with the restoration of mtDNA and loss of the acquired adaptation (Fig. 3E, figs. S5C, D).

The above observations strongly suggest that the loss of OXPHOS activity was the predominant mechanism responsible for the swiftness of the first adaptation phase. Considering that 68% of the realized adaptation was achieved during this phase, the major fraction of paraquat-induced O_2_^•−^ production is therefore arguably associated with the OXPHOS system (*19*).

### The deletion of mtDNA segments requires *SOD2*

Based on experiments that led us to conclude that hydrogen peroxide (H_2_O_2_) or its downstream degradation products did not play a prominent causative role in inducing the deletion of mtDNA segments (figs. S6A, B, supplementary text 4), we hypothesized that mitochondrial O_2_^•−^ was directly responsible for this. The superoxide dismutases Sod1 (Cu/ZnSOD) and Sod2 (MnSOD), located in the cytosol and mitochondrial intermembrane space and in the mitochondrial matrix, respectively, have both previously been linked to signaling responses associated with O_2_^•−^ (*20, 27*). We therefore probed how *sod1Δ* and *sod2Δ* cells responded to paraquat compared to the wild type. While the *sod1Δ* populations adapted as wild type cells, the *sod2Δ* populations mostly displayed no reduction in cell doubling time over 10 growth cycles when exposed to the same stress levels (Figs. 4A, B, fig. S6C, supplementary text 5).

**Figure 4.**
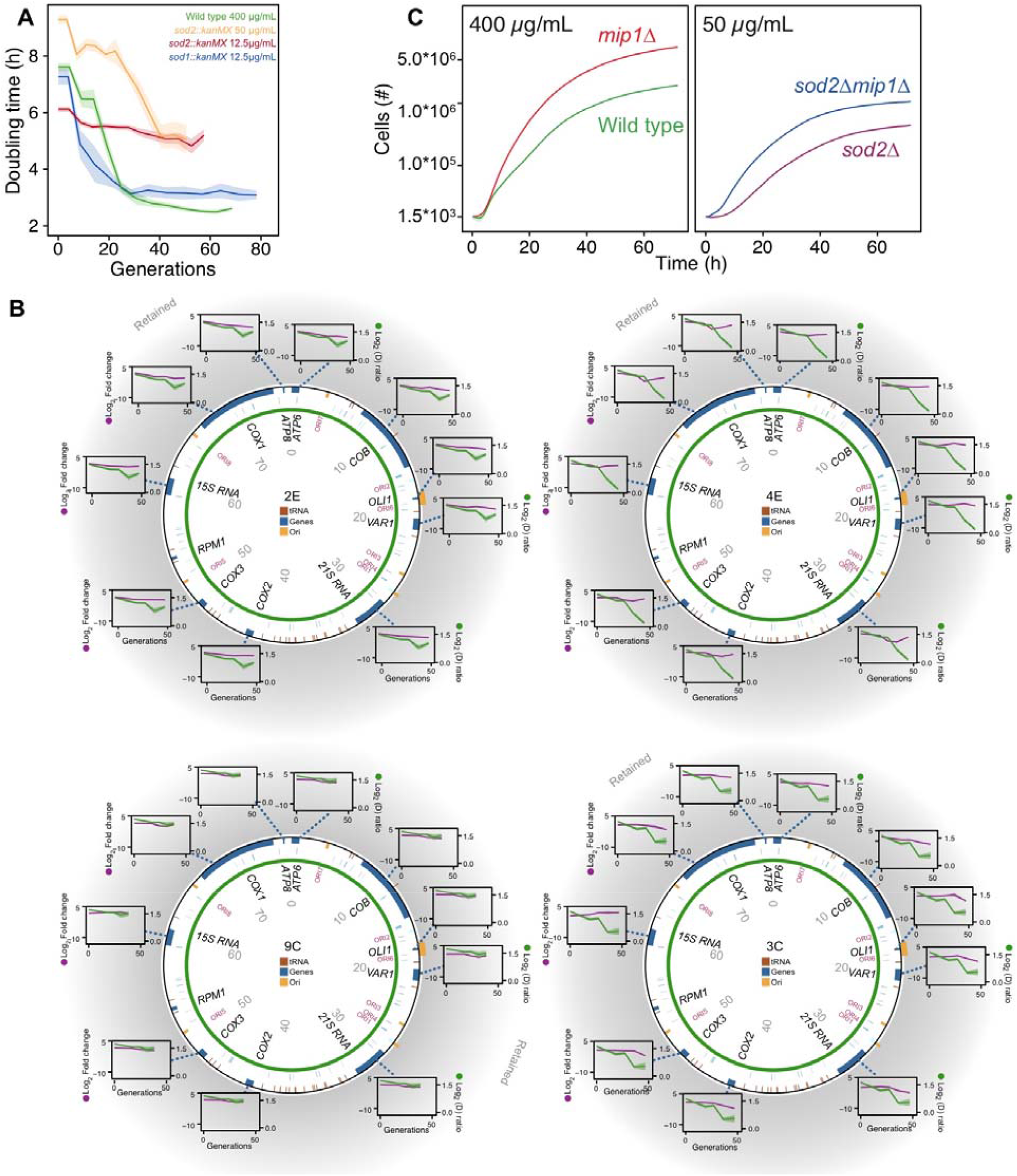
The mtDNA editing critically involves Sod2. (A) Doubling time (*h*) in wild type (400 µg/mL paraquat: green), *sod2Δ* (12.5 µg/mL paraquat: red; 50 µg/mL paraquat: yellow) and *sod1Δ* (12.5 µg/mL paraquat: blue) cell populations adapting to equivalent stress levels. (**B**) mtDNA change in *sod2Δ* cell populations 2E, 4E, 9C and 3C adapting to 12.5 µg/mL paraquat. *Circle:* mtDNA (77 kb) before exposure to paraquat. Genes, origins of replication and nucleotide positions (kb) are indicated. *Colored fields:* mtDNA deletions with concerted copy number change. *Diagrams:* mtDNA copy number change (left *y*-axis, purple line, (*n=*2) of individual mtDNA genes during the adaptation (right *y*-axis, green line). Shade: S.E.M. (**C**) Mean growth of wild type (*n=*480; green), *mip1Δ* (*n=*288; red), *sod2Δ* (*n=*96; purple) and *sod2Δmip1Δ* (*n=*384; blue) cell populations in the presence of 400 µg/mL (left) and 50 µg/mL (right) paraquat.

Using qPCR, we then tracked the pattern of mtDNA deletions in eight *sod2Δ* populations exposed to paraquat. In four of the populations, the copy numbers of all mtDNA genes were retained at or near pre-stress level (Fig. 4B). Two of these failed to adapt to paraquat, while the other two showed a weak and a much-delayed reduction in cell doubling time relative to the wild type, indicating selection for small-effect mutations not associated with mtDNA deletion. The remaining four populations showed a much-delayed reduction in cell doubling time that coincided with the acquisition of a single mtDNA deletion (fig. S6D). To clarify the importance of mtDNA deletions in relation to the Sod2-dependency, we deleted Mip1 in a *sod2Δ* background, and found that the induced loss of mtDNA increased growth as in wild type cells (Fig. 4C). This suggests that the delayed adaptation to intramitochondrial O _2_^•−^ stress in cells lacking Sod2 is because the mtDNA deletions emerge at a much lower rate. Taken together, the above results strongly suggest that Sod2 is a key factor in the causal chain of events leading to the mtDNA deletions underlying the initial rapid adaptation to paraquat. As the data clearly show that the deletion of mtDNA segments causing loss of OXPHOS activity is under regulatory control, we think it is apt to denote this process as ‘mtDNA editing’ (*28*).

### The mtDNA editing process requires anterograde mito-nuclear communication

The above results led us to suspect that the underlying regulatory circuitry might involve mitochondrial-nuclear communication. In budding yeast, deletion of mtDNA alters the expression of a multitude of genes resulting in increased glycolytic production of ATP (*29*), dubbed the retrograde response. Upon mitochondrial OXPHOS dysfunction, the cytosolic protein Rtg2 (*30*) causes the transcription factors Rtg1 and Rtg3 to translocate from the cytosol to the nucleus where they together activate retrograde transcription (*31, 32*). As this mechanism is the most well-documented channel in yeast for communicating mitochondrial dysfunction to the nucleus, and in particular mtDNA deletion (*33*), we exposed *rtg2Δ* and *rtg3Δ* cell populations (*n=*16) to paraquat for 80 generations. In both cases, all cell populations failed to adapt (Fig. 5A), and their capacity for respiratory growth was virtually unperturbed at the end of the experiment (Fig. 5B). This indicates that the two proteins induce anterograde mito-nuclear communication critical for activation of the mtDNA editing process.

**Figure 5.**
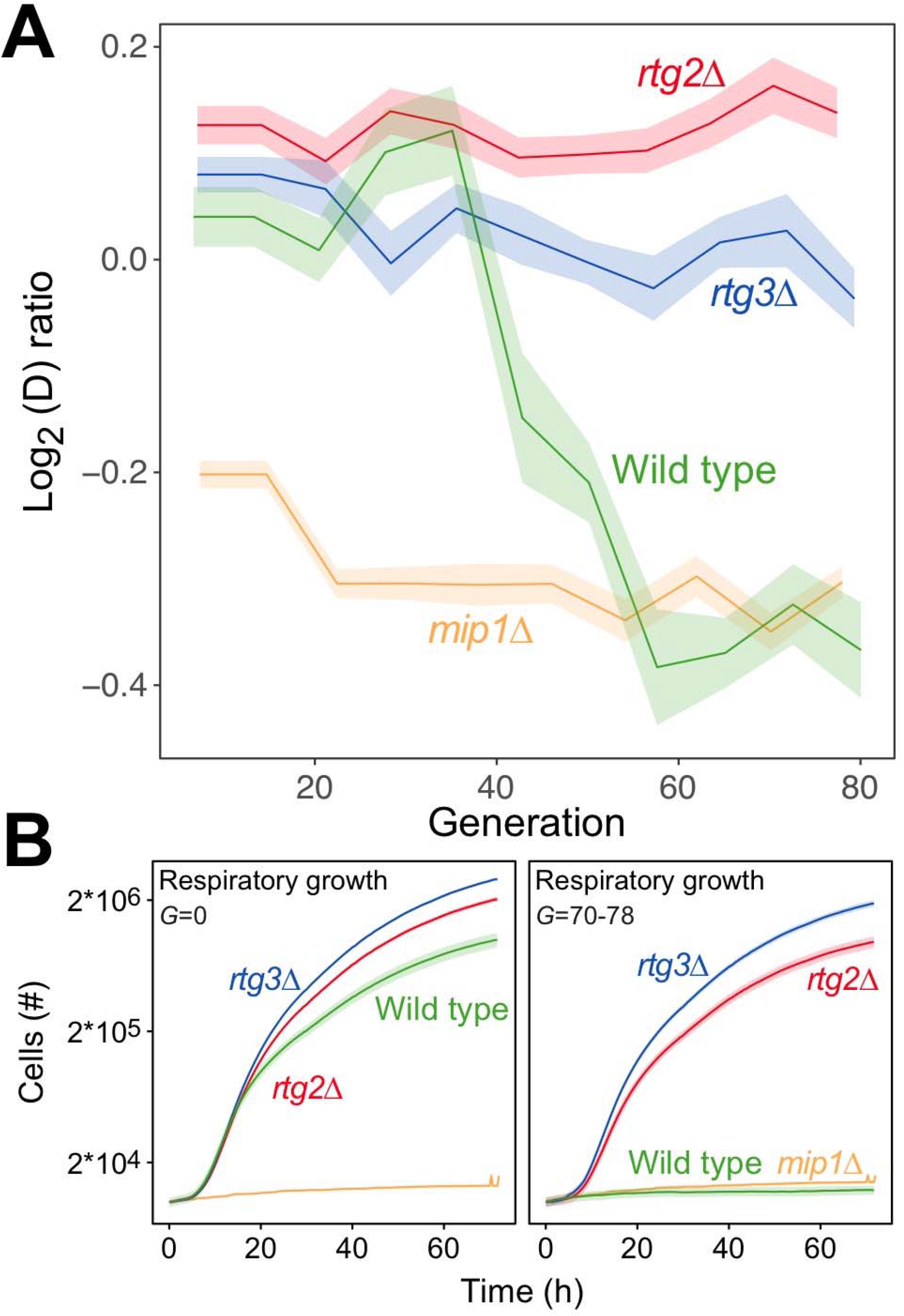
The mtDNA editing critically involves anterograde mito-nuclear communication. (**A**) Doubling time adaptation of 18 wild type, *rtg2Δ, rtg3Δ* and *mip1Δ* cell populations to 400 µg/mL of paraquat. Shade: S.E.M. Each population type measured at *n*=4. (**B**) Respiratory (glycerol) growth of wild type, *rtg2Δ, rtg3Δ* and *mip1Δ* cell populations, before (left) and after (right) 70-78 generations of paraquat adaptation. Shade: S.E.M (*n*=72-144 populations, each measured at *n*=1).

### Sustained mtDNA deletion causes irrevocable mitochondrial impairment

We found that 44 of the 96 populations had completely lost their capacity for restoring the pool of intact mtDNA genomes back to pre-stress levels after 24 generations of paraquat exposure, and after 242 generations all did (fig. S7). Moreover, in the five sequenced cell populations, the capacity to restore the copy numbers of intact mtDNA and the respiratory growth after removal of stress was lost between 15 to 42 generations of paraquat exposure (Fig. 6A). This suggest that the deletion of mtDNA genes continued after the first adaptation phase in a minor fraction of cells without providing a recognizable further reduction in cell doubling time.

**Figure 6.**
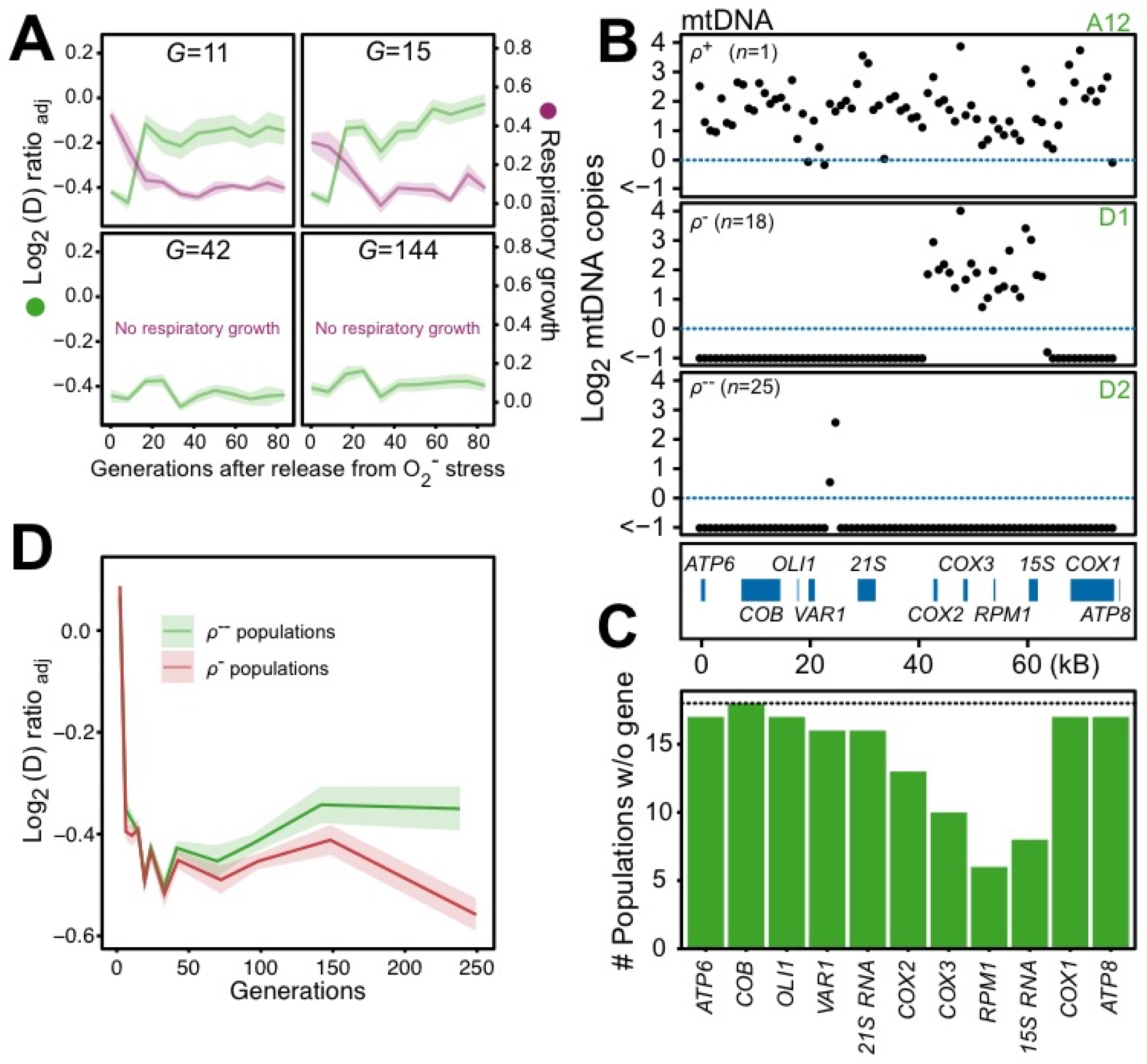
Chronic exposure to paraquat causes irreversible mitochondrial impairment by sustained mtDNA editing. (**A**) Paraquat adapting cell populations (*G* = generations of exposure to paraquat) ultimately lose their capacity to recover respiratory (glycerol) growth (right *y*-axis, purple line, log_2_ doubling time relative to founder) and the loss coincides with the genetic fixation of the paraquat adaptation (left *y*-axis, green line). Shade: S.E.M. of 5 populations, each measured at *n=*5. (**B**) All but one (*ρ*^+^) sequenced cell population adapted to long-term paraquat stress (*t*_*50*_) retain only small (6-30 kb; *ρ*^−^) or very small (<2 kb, *ρ*^--^) mtDNA segments. *Panels:* Representative populations. *y*-axis: mtDNA copy number (median coverage in 1 kb windows relative to haploid nuclear genome). Gene positions are indicated. (**C**) Number of *ρ*^−^ populations after long-term paraquat exposure (*t*_*50*_) in which the specified mtDNA gene was lost. (**D**) The *ρ*^--^ populations became less fit than the *ρ*^−^ populations during a long-term exposure to paraquat.

The sustentation of mtDNA deletions led us to suspect that a long-term O _2_^•−^ stress would finally lead to a complete loss of intact mtDNA genomes. We therefore assayed the mtDNA loss in 44 sequenced endpoint (*t*_*50*_) populations. Out of these, 25 populations had almost completely lost their entire 77 kb mtDNA (97-99%) (fig S8). 18 endpoint populations remained in a rho negative (*ρ*^−^) state where they still possessed 6 to 34 kb mtDNA segments with copy numbers somewhat above the original founder levels (mean: 20% increase). To assess whether there was an adaptive advantage associated with the sustained depletion of mtDNA, we compared the doubling times of the *ρ*^−^ populations still retaining 6 to 34 kb mtDNA segments with those that had lost almost all their mtDNA. Intriguingly, the latter group consistently grew slower on paraquat than the former (Fig. 6D). This suggests that the sustained mtDNA depletion under long-term stress, leading to irrevocable loss of intact mtDNA genomes, is a maladaptive response driven by prolonged induction of a regulatory mechanism dimensioned by natural selection to handle O_2_^•−^ stress it can successfully deal with before the pool of intact mtDNA genomes becomes annihilated.

### Chromosome duplications explain the second adaptation phase

*mip1Δ* cells lacking mtDNA were not fully tolerant to paraquat (Fig. S9A), consistent with that wild type cells realized 23.3% of their adaptation potential in a second adaptation phase (Fig. 1A). The much slower doubling time reduction characterizing this second phase suggested that in this case the adaptation to on- or off-target effects of paraquat could be explained by a Darwinian mutation/selection process (fig. S2). Analysis of the sequence data of 44 random endpoint (*t*_*50*_) populations led us to conclude that nuclear point mutations were unlikely to have played a prominent causative role, though (figs. S9B-C). However, duplications of chromosome II, III and V were common, and we found that these were able to explain the reduction in cell doubling time during the second adaptation phase (fig S10 and supplementary text 6).

## Discussion

The adaptation patterns of the *sod2Δ, mip1Δ, rtg2Δ* and *rtg3Δ* populations relative to the wild type are hard to reconcile with the operation of random O_2_^•−^ induced mtDNA deletions (e.g. caused by genomic instability) and subsequent selection of cells possessing OXPHOS-impaired mitochondria. However, the specificity of the mtDNA deletion mechanism in terms of which mtDNA segments are lost in the first adaptation phase does seem to be moderate. Still, there was a clear preference for deletions within the *COX1-VAR1* region, and the segment containing *COX3, RPM1* and 15S RNA were not deleted in this phase. The fact that copy number homeostasis was observed not just for these three genes, but also for genes in undeleted segments within the *COX1-VAR1* region, indicates that partial deletion of genes within this region was sufficient for providing the marked reduction in cell doubling time during the first adaptation phase. This interpretation makes sense from an evolutionary point of view: natural selection would not be able to increase the specificity of the mtDNA editing program beyond the point where no further adaptation is achieved. However, the data suggest that even the moderate specificity characterizing the early adaptation phase is lost by sustained operation of the mtDNA deletion mechanism during long-term oxidative stress.

The restoration of intact mtDNA genomes back to pre-stress levels after release from paraquat during the first adaptation phase (Fig. 3E) is almost complete after just 2-3 growth cycles despite that there is no apparent selective advantage attached (Figs. 1B, C). This signifies that it is under strict regulatory control. A possible mechanism is selective fusion of mitochondria with intact mtDNA genomes (*34, 35*) during the swift return to tubular morphology after cessation of stress (Fig. 2B), and subsequent removal of mitochondria with OXPHOS deprived genomes by mitophagy. Such selective fusion may also be an integral part of the fission/fusion dynamics such that cells achieve the clearance of non-intact mtDNA genomes only after several fission/fusion cycles. In this case, one would expect the restoration to be under the control of the constituent homeostatic regulatory machinery. If the associated feedback control mechanism is oblivious to non-intact mtDNA genomes (*36*), and the restoration is faster than the removal of the OXPHOS-deprived genomes, one would expect a temporal overshoot of mtDNA copy numbers before a return to prestress levels. Our data are in clear agreement with this notion (fig. S5D).

Besides providing extensive support to the conception that the disclosed mtDNA-editing system defines a regulatory layer between the primary antioxidant defenses and canonical mitophagy for coping with intramitochondrial supraphysiological O_2_^•−^ challenges, our data bring fresh perspectives to the table concerning (i) the relationship between stress-induced mitochondrial fragmentation and canonical mitophagy (*37*), (ii) under which conditions do mitochondria deprived of OXPHOS genes produce more O_2_^•−^ due to increased electron leakage (*38, 39*), (iii) whether clonal expansion is the main mechanism underlying the propagation of mitochondria containing deletions (*40*), and (iv) under which conditions, and to which degree, does the main retrograde response to mtDNA deletion in yeast mediate two-way mito-nuclear communication (*33*). They also support the emerging notion that selective mitophagy is an important mechanism for local mitochondrial repair (*13*), and that selective mitophagy may be deliberately repressed while the cell experiences O_2_^•−^ stress.

Available data on the cancer therapeutics doxorubicin and cisplatin suggest that our main results may extend to post-mitotic cells. They both accumulate in mitochondria (*41, 42*), where they induce O_2_^•−^ production through redox cycling (*43, 44*). Their long-term administration cause oxidative injury to a variety of cells and tissues (*44*–*47*), and they both increase the frequency of mtDNA deletions (*42, 48*). Arrestment of OXPHOS activity by mtDNA editing, followed by homeostatic restoration of the pool of intact mitochondrial genomes, may thus be a generic eukaryotic adaptation to obviate a more costly mitophagic response in a variety of situations. If so, the biomedical implications in terms of the etiology of age-related disease and therapeutic opportunities appear to be noteworthy.

## Acknowledgement

We thank Johan Hallin and Lars-Göran Ottosson for help and advice with strain construction and design of adaptation experiment, Olga Kourtchenko for help with designing nitrogen-limited environments, and Tom Kirkwood for instrumental comments to an earlier version of this paper. The authors acknowledge Illumina sequencing technical support from the Science for Life Laboratory (SciLife), the National Genomics Infrastructure, NGI and National Bioinformatics Infrastructure in Sweden and PacBio sequencing technical support from the Norwegian Sequencing Centre. We acknowledge the Centre for Cellular Imaging at the University of Gothenburg and the National Microscopy Infrastructure (VR-RFI 2016-00968) for assistance with the confocal microscopy.

## Funding

Swedish Research Council (2014-6547, 2014-4605, 2015-05427, 2018-03638 and 2018-03453) to JW, MM and JLH

Cancerfonden (2017-778) to MM

Research Council of Norway (178901/V30 and 222364/F20) to SWO

Agence Nationale de la Recherche (ANR-11-LABX-0028-01, ANR-13-BSV6-0006-01, ANR-15-IDEX-01, ANR-16-CE12-0019 and ANR-18-CE12-0004) to GL

## Author contributions

Conceptualization: SWO, JW, MM, ABG and GL

Methodology: JW, SWO, ABG, SS, GL, LL, JLH and JL

Formal analysis: SS, CGP, KP, CG, PG, EDH, JL, JXY, CG, LL, MZ, ABG, TÄ

Investigation: SS, CGP, KP, CG, PG, EDH, JL, JXY, CG, LL

Writing – original draft: SS, SWO, JW, MM

Writing – review and editing: all authors

Visualization: SS, JW, KP, JL, JXY, CG

Supervision: JW, SWO, GL, MM, JLH

Project administration: JW, SWO, ABG

Funding acquisition: JW, SWO, ABG, GL, MM, JLH

## Data and materials availability

Sequence data that support the findings of this study have been deposited in Sequencing Read Archive (SRA) with the accession codes PRJNA622836. The growth phenotyping code can be found at https://github.com/Scan-o-Matic/scanomatic.git, the simulation code at https://github.com/HelstVadsom/GenomeAdaptation.git and the imaging code at https://github.com/CamachoDejay/SStenberg_3Dyeast_tools. The authors declare that all other data supporting the findings of this study are available within the paper as Supplemental Information Data S1-S30, which can be previewed at https://data.mendeley.com/datasets/mvx7t7rw2d/draft?a=95381e47-dc80-47af-85ab-e0478912a209. All unique strains and stored populations generated in this study are available from the Lead Contact without restriction.

## Supplementary Materials

Materials and Methods

Supplementary Text S1 to S6

Figs. S1 to S10

Table S1

References 49-77

Data S1 to S30

## References

1. N. Sun, R. J. Youle, T. Finkel, The Mitochondrial Basis of Aging. Mol. Cell. 61, 654–666 (2016).

2. H. Hu, C.-C. Tan, L. Tan, J.-T. Yu, A Mitocentric View of Alzheimer’s Disease. Mol. Neurobiol. 54, 6046–6060 (2017).

3. N. Ammal Kaidery, B. Thomas, Current perspective of mitochondrial biology in Parkinson’s disease. Neurochem. Int. 117, 91–113 (2018).

4. R. T. Hepple, Impact of aging on mitochondrial function in cardiac and skeletal muscle. Free Radic. Biol. Med. 98, 177–186 (2016).

5. J. M. T. Hyttinen, J. Viiri, K. Kaarniranta, J. Błasiak, Mitochondrial quality control in AMD: does mitophagy play a pivotal role? Cell. Mol. Life Sci. 75, 2991–3008 (2018).

6. K. J. Krishnan, A. K. Reeve, D. C. Samuels, P. F. Chinnery, J. K. Blackwood, R. W. Taylor, S. Wanrooij, J. N. Spelbrink, R. N. Lightowlers, D. M. Turnbull, What causes mitochondrial DNA deletions in human cells? Nat. Genet. 40, 275–279 (2008).

7. N. Nissanka, M. Minczuk, C. T. Moraes, Mechanisms of Mitochondrial DNA Deletion Formation. Trends Genet. 35, 235–244 (2019).

8. G. A. Fontana, H. L. Gahlon, Mechanisms of replication and repair in mitochondrial DNA deletion formation. Nucleic Acids Res. 48, 11244–11258 (2020).

9. J. J. Lemasters, Selective Mitochondrial Autophagy, or Mitophagy, as a Targeted Defense Against Oxidative Stress, Mitochondrial Dysfunction, and Aging. Rejuvenation Res. 8, 3–5 (2005).

10. A. S. Bess, T. L. Crocker, I. T. Ryde, J. N. Meyer, Mitochondrial dynamics and autophagy aid in removal of persistent mitochondrial DNA damage in Caenorhabditis elegans. Nucleic Acids Res. 40, 7916–7931 (2012).

11. K. Palikaras, N. Tavernarakis, Mitochondrial homeostasis: the interplay between mitophagy and mitochondrial biogenesis. Exp. Gerontol. 56, 182–8 (2014).

12. L. Sedlackova, V. I. Korolchuk, Mitochondrial quality control as a key determinant of cell survival. Biochim. Biophys. Acta - Mol. Cell Res. 1866, 575–587 (2019).

13. Å. B. Gustafsson, G. W. Dorn, Evolving and expanding the roles of mitophagy as a homeostatic and pathogenic process. Physiol. Rev. 99, 853–892 (2019).

14. H. Sies, C. Berndt, D. P. Jones, Oxidative Stress. Annu. Rev. Biochem. 86, 715–748 (2017).

15. T. Shpilka, C. M. Haynes, The mitochondrial UPR: Mechanisms, physiological functions and implications in ageing. Nat. Rev. Mol. Cell Biol. 19, 109–120 (2018).

16. M. Y. W. Ng, T. Wai, A. Simonsen, Quality control of the mitochondrion. Dev. Cell. 56, 881–905 (2021).

17. M. De Chiara, B. Barré, K. Persson, A. O. Chioma, A. Irizar, J. Schacherer, J. Warringer, G. Liti, bioRxiv, in press, doi:10.1101/2020.02.08.939314.

18. H. M. Cochemé, M. P. Murphy, Complex I is the major site of mitochondrial superoxide production by paraquat. J. Biol. Chem. 283, 1786–1798 (2008).

19. P. R. Castello, D. A. Drechsel, M. Patel, Mitochondria are a major source of paraquat-induced reactive oxygen species production in the brain. J. Biol. Chem. 282, 14186–93 (2007).

20. X. Zou, B. A. Ratti, J. G. O’Brien, S. O. Lautenschlager, D. R. Gius, M. G. Bonini, Y. Zhu, Manganese superoxide dismutase (SOD2): is there a center in the universe of mitochondrial redox signaling? J. Bioenerg. Biomembr. 49, 325–333 (2017).

21. A. B. Gjuvsland, E. Zörgö, J. K. Samy, S. Stenberg, I. H. Demirsoy, F. Roque, E. Maciaszczyk-Dziubinska, M. Migocka, E. Alonso-Perez, M. Zackrisson, R. Wysocki, M. J. Tamás, I. Jonassen, S. W. Omholt, J. Warringer, Disentangling genetic and epigenetic determinants of ultrafast adaptation. Mol. Syst. Biol. 12, 892 (2016).

22. H. G. Sprenger, T. Langer, The Good and the Bad of Mitochondrial Breakups. Trends Cell Biol. 29, 888–900 (2019).

23. M. Frank, S. Duvezin-Caubet, S. Koob, A. Occhipinti, R. Jagasia, A. Petcherski, M. O. Ruonala, M. Priault, B. Salin, A. S. Reichert, Mitophagy is triggered by mild oxidative stress in a mitochondrial fission dependent manner. Biochim. Biophys. Acta - Mol. Cell Res. 1823, 2297–2310 (2012).

24. C. H.-L. Hung, S. S.-Y. Cheng, Y.-T. Cheung, S. Wuwongse, N. Q. Zhang, Y.-S. Ho, S. M.-Y. Lee, R. C.-C. Chang, A reciprocal relationship between reactive oxygen species and mitochondrial dynamics in neurodegeneration. Redox Biol. 14, 7–19 (2018).

25. Y. Liu, K. Okamoto, Regulatory mechanisms of mitophagy in yeast. Biochim. Biophys. Acta - Gen. Subj. 1865, 129858 (2021).

26. T. Lodi, C. Dallabona, C. Nolli, P. Goffrini, C. Donnini, E. Baruffini, DNA polymerase Î^3^and disease: what we have learned from yeast. Front. Genet. 6 (2015), doi:10.3389/fgene.2015.00106.

27. A. R. Reddi, V. C. Culotta, SOD1 integrates signals from oxygen and glucose to repress respiration. Cell. 152, 224–35 (2013).

28. Merriam-Webster, Edit. Merriam-Webster’s Coll. Thes., (available at https://unabridged.merriam-webster.com/thesaurus/edit).

29. C. B. Epstein, J. A. Waddle, W. Hale IV, V. Davé, J. Thornton, T. L. Macatee, H. R. Garner, R. A. Butow, Genome-wide responses to mitochondrial dysfunction. Mol. Biol. Cell. 12, 297–308 (2001).

30. T. Sekito, J. Thornton, R. A. Butow, Mitochondria-to-Nuclear Signaling Is Regulated by the Subcellular Localization of the Transcription Factors Rtg1p and Rtg3p. Mol. Biol. Cell. 11, 2103–2115 (2000).

31. B. A. Rothermel, A. W. Shyjan, J. L. Etheredge, R. A. Butow, Transactivation by Rtg1p, a Basic Helix-Loop-Helix Protein That Functions in Communication between Mitochondria and the Nucleus in Yeast. J. Biol. Chem. 270, 29476–29482 (1995).

32. B. A. Rothermel, J. L. Thornton, R. A. Butow, Rtg3p, a Basic Helix-Loop-Helix/Leucine Zipper Protein that Functions in Mitochondrial-induced Changes in Gene Expression, Contains Independent Activation Domains. J. Biol. Chem. 272, 19801–19807 (1997).

33. N. Guaragnella, L. P. Coyne, X. J. Chen, S. Giannattasio, Mitochondria–cytosol–nucleus crosstalk: learning from Saccharomyces cerevisiae. FEMS Yeast Res. 18 (2018), doi:10.1093/femsyr/foy088.

34. G. Twig, A. Elorza, A. J. A. Molina, H. Mohamed, J. D. Wikstrom, G. Walzer, L. Stiles, S. E. Haigh, S. Katz, G. Las, J. Alroy, M. Wu, B. F. Py, J. Yuan, J. T. Deeney, B. E. Corkey, O. S. Shirihai, Fission and selective fusion govern mitochondrial segregation and elimination by autophagy. EMBO J. 27, 433–446 (2008).

35. T. Ban, T. Ishihara, H. Kohno, S. Saita, A. Ichimura, K. Maenaka, T. Oka, K. Mihara, N. Ishihara, Molecular basis of selective mitochondrial fusion by heterotypic action between OPA1 and cardiolipin. Nat. Cell Biol. 19, 856–863 (2017).

36. A. Kowald, T. B. L. Kirkwood, Transcription could be the key to the selection advantage of mitochondrial deletion mutants in aging. Proc. Natl. Acad. Sci. 111, 2972–2977 (2014).

37. D. Zorov, I. Vorobjev, V. Popkov, V. Babenko, L. Zorova, I. Pevzner, D. Silachev, S. Zorov, N. Andrianova, E. Plotnikov, Lessons from the Discovery of Mitochondrial Fragmentation (Fission): A Review and Update. Cells. 8, 175 (2019).

38. L. Aerts, V. A. Morais, in Parkinson’s Disease (Elsevier, 2017; https://linkinghub.elsevier.com/retrieve/pii/B978012803783600002X), pp. 41–75.

39. S. Sabnam, H. Rizwan, S. Pal, A. Pal, CEES-induced ROS accumulation regulates mitochondrial complications and inflammatory response in keratinocytes. Chem. Biol. Interact. 321, 109031 (2020).

40. G. S. Nido, C. Dölle, I. Flønes, H. A. Tuppen, G. Alves, O. B. Tysnes, K. Haugarvoll, C. Tzoulis, Ultradeep mapping of neuronal mitochondrial deletions in Parkinson’s disease. Neurobiol. Aging. 63, 120–127 (2018).

41. B. Kalyanaraman, J. Joseph, S. Kalivendi, S. Wang, E. Konorev, S. Kotamraju, Doxorubicin-induced apoptosis: Implications in cardiotoxicity. Mol. Cell. Biochem. 234–235, 119–124 (2002).

42. G. Genc, A. Okuyucu, B. C. Meydan, O. Yavuz, O. Nisbet, M. Hokelek, A. Bedir, O. Ozkaya, Effect of free creatine therapy on cisplatin-induced renal damage. Ren. Fail. 36, 1108–1113 (2014).

43. S. S. Malhi, A. Budhiraja, S. Arora, K. R. Chaudhari, K. Nepali, R. Kumar, H. Sohi, R. S. R. Murthy, Intracellular delivery of redox cycler-doxorubicin to the mitochondria of cancer cell by folate receptor targeted mitocancerotropic liposomes. Int. J. Pharm. 432, 63–74 (2012).

44. Z. Song, H. Chang, N. Han, Z. Liu, Y. Liu, H. Wang, J. Shao, Z. Wang, H. Gao, J. Yin, He-Wei granules (HWKL) combat cisplatin-induced nephrotoxicity and myelosuppression in rats by inhibiting oxidative stress, inflammatory cytokines and apoptosis. RSC Adv. 7, 19794–19807 (2017).

45. M. Songbo, H. Lang, C. Xinyong, X. Bin, Z. Ping, S. Liang, Oxidative stress injury in doxorubicin-induced cardiotoxicity. Toxicol. Lett. 307, 41–48 (2019).

46. J. F. Moruno-Manchon, N. Uzor, S. R. Kesler, J. S. Wefel, D. M. Townley, A. S. Nagaraja, S. Pradeep, L. S. Mangala, A. K. Sood, A. S. Tsvetkov, Peroxisomes contribute to oxidative stress in neurons during doxorubicin-based chemotherapy. Mol. Cell. Neurosci. 86, 65–71 (2018).

47. X. Ren, J. T. R. Keeney, S. Miriyala, T. Noel, D. K. Powell, L. Chaiswing, S. Bondada, D. K. St. Clair, D. A. Butterfield, The triangle of death of neurons: Oxidative damage, mitochondrial dysfunction, and loss of choline-containing biomolecules in brains of mice treated with doxorubicin. Advanced insights into mechanisms of chemotherapy induced cognitive impairment (“chemobr. Free Radic. Biol. Med. 134, 1–8 (2019).

48. K. Adachi, Y. Fujiura, F. Mayumi, A. Nozuhara, Y. Sugiu, T. Sakanashi, T. Hidaka, H. Toshima, A Deletion of Mitochondrial DNA in Murine Doxorubicin-Induced Cardiotoxicity. Biochem. Biophys. Res. Commun. 195, 945–951 (1993).

49. G. Liti, D. M. Carter, A. M. Moses, J. Warringer, L. Parts, S. A. James, R. P. Davey, I. N. Roberts, Burt, V. Koufopanou, I. J. Tsai, C. M. Bergman, D. Bensasson, M. J. T. O’Kelly, A. van Oudenaarden, D. B. H. Barton, E. Bailes, A. N. Nguyen, M. Jones, M. A. Quail, I. Goodhead, S. Sims, F. Smith, A. Blomberg, R. Durbin, E. J. Louis, Population genomics of domestic and wild yeasts. Nature. 458, 337–41 (2009).

50. M. Gaisne, A. M. Bécam, J. Verdière, C. J. Herbert, A “natural” mutation in Saccharomyces cerevisiae strains derived from S288c affects the complex regulatory gene HAP1 (CYP1). Curr. Genet. 36, 195–200 (1999).

51. J. Zhu, Z.-T. Zhang, S.-W. Tang, B.-S. Zhao, H. Li, J.-Z. Song, D. Li, Z. Xie, A Validated Set of Fluorescent-Protein-Based Markers for Major Organelles in Yeast (Saccharomyces cerevisiae). MBio. 10, 1–19 (2019).

52. A. Gutiérrez, M. Sancho, G. Beltran, J. M. Guillamon, J. Warringer, Replenishment and mobilization of intracellular nitrogen pools decouples wine yeast nitrogen uptake from growth. Appl. Microbiol. Biotechnol. 100, 3255–3265 (2016).

53. M. Zackrisson, J. Hallin, L.-G. Ottosson, P. Dahl, E. Fernandez-Parada, E. Ländström, L. Fernandez-Ricaud, P. Kaferle, A. Skyman, S. Stenberg, S. Omholt, U. Petrovič, J. Warringer, A. Blomberg, Scan-o-matic: High-Resolution Microbial Phenomics at a Massive Scale. G3 (Bethesda). 6, 3003–14 (2016).

54. S. W. Wingett, S. Andrews, FastQ Screen: A tool for multi-genome mapping and quality control. F1000Research. 7, 1338 (2018).

55. A. Dobin, C. A. Davis, F. Schlesinger, J. Drenkow, C. Zaleski, S. Jha, P. Batut, M. Chaisson, T. R. Gingeras, STAR: ultrafast universal RNA-seq aligner. Bioinformatics. 29, 15–21 (2013).

56. Y. Liao, G. K. Smyth, W. Shi, featureCounts: an efficient general purpose program for assigning sequence reads to genomic features. Bioinformatics. 30, 923–930 (2014).

57. M. I. Love, W. Huber, S. Anders, Moderated estimation of fold change and dispersion for RNA-seq data with DESeq2. Genome Biol. 15, 550 (2014).

58. J.-X. Yue, J. Li, L. Aigrain, J. Hallin, K. Persson, K. Oliver, A. Bergström, P. Coupland, J. Warringer, M. C. Lagomarsino, G. Fischer, R. Durbin, G. Liti, Contrasting evolutionary genome dynamics between domesticated and wild yeasts. Nat. Genet. 49, 913–924 (2017).

59. J.-X. Yue, G. Liti, Long-read sequencing data analysis for yeasts. Nat. Protoc. 13, 1213–1231 (2018).

60. G. Giaever, A. M. Chu, L. Ni, C. Connelly, L. Riles, S. Véronneau, S. Dow, A. Lucau-Danila, K. Anderson, B. André, A. P. Arkin, A. Astromoff, M. El-Bakkoury, R. Bangham, R. Benito, S. Brachat, S. Campanaro, M. Curtiss, K. Davis, A. Deutschbauer, K.-D. Entian, P. Flaherty, F. Foury, D. J. Garfinkel, M. Gerstein, D. Gotte, U. Güldener, J. H. Hegemann, S. Hempel, Z. Herman, D. F. Jaramillo, D. E. Kelly, S. L. Kelly, P. Kötter, D. LaBonte, D. C. Lamb, N. Lan, H. Liang, H. Liao, L. Liu, C. Luo, M. Lussier, R. Mao, P. Menard, S. L. Ooi, J. L. Revuelta, C. J. Roberts, M. Rose, P. Ross-Macdonald, B. Scherens, G. Schimmack, B. Shafer, D. D. Shoemaker, S. Sookhai-Mahadeo, R. K. Storms, J. N. Strathern, G. Valle, M. Voet, G. Volckaert, C. Wang, T. R. Ward, J. Wilhelmy, E. A. Winzeler, Y. Yang, G. Yen, E. Youngman, K. Yu, H. Bussey, J. D. Boeke, M. Snyder, P. Philippsen, R. W. Davis, M. Johnston, Functional profiling of the Saccharomyces cerevisiae genome. Nature. 418, 387–91 (2002).

61. D. C. Zebrowski, D. B. Kaback, A simple method for isolating disomic strains of Saccharomyces cerevisiae. Yeast. 25, 321–6 (2008).

62. Y. O. Zhu, M. L. Siegal, D. W. Hall, D. A. Petrov, Precise estimates of mutation rate and spectrum in yeast. Proc. Natl. Acad. Sci. U. S. A. 111, E2310–8 (2014).

63. G. Chevereau, M. Dravecká, T. Batur, A. Guvenek, D. H. Ayhan, E. Toprak, T. Bollenbach, Quantifying the Determinants of Evolutionary Dynamics Leading to Drug Resistance. PLoS Biol. 13, e1002299 (2015).

64. R. Vaser, S. Adusumalli, S. N. Leng, M. Sikic, P. C. Ng, SIFT missense predictions for genomes. Nat. Protoc. 11, 1–9 (2016).

65. H.-H. Chou, H.-C. Chiu, N. F. Delaney, D. Segrè, C. J. Marx, Diminishing returns epistasis among beneficial mutations decelerates adaptation. Science. 332, 1190–2 (2011).

66. A. I. Khan, D. M. Dinh, D. Schneider, R. E. Lenski, T. F. Cooper, Negative epistasis between beneficial mutations in an evolving bacterial population. Science. 332, 1193–6 (2011).

67. P. Hawes, C. L. Netherton, M. Mueller, T. Wileman, P. Monaghan, Rapid freeze-substitution preserves membranes in high-pressure frozen tissue culture cells. J. Microsc. 226, 182–9 (2007).

68. E. S. Reynolds, The use of lead citrate at high pH as an electron-opaque stain in electron microscopy. J. Cell Biol. 17, 208–12 (1963).

69. J. R. Kremer, D. N. Mastronarde, J. R. McIntosh, Computer visualization of three-dimensional image data using IMOD. J. Struct. Biol. 116, 71–6 (1996).

70. H. G. Crabtree, Observations on the carbohydrate metabolism of tumours. Biochem. J. 23, 536–45 (1929).

71. H. Sies, D. P. Jones, Reactive oxygen species (ROS) as pleiotropic physiological signalling agents. Nat. Rev. Mol. Cell Biol. 21, 363–383 (2020).

72. K. Bodvard, K. Peeters, F. Roger, N. Romanov, A. Igbaria, N. Welkenhuysen, G. Palais, W. Reiter, M. B. Toledano, M. Käll, M. Molin, Light-sensing via hydrogen peroxide and a peroxiredoxin. Nat. Commun. 8, 14791 (2017).

73. S. Okazaki, T. Tachibana, A. Naganuma, N. Mano, S. Kuge, Multistep Disulfide Bond Formation in Yap1 Is Required for Sensing and Transduction of H2O2 Stress Signal. Mol. Cell. 27, 675–688 (2007).

74. A. Delaunay, D. Pflieger, M.-B. Barrault, J. Vinh, M. B. Toledano, A Thiol Peroxidase Is an H2O2 Receptor and Redox-Transducer in Gene Activation. Cell. 111, 471–481 (2002).

75. A. Hori, M. Yoshida, T. Shibata, F. Ling, Reactive oxygen species regulate DNA copy number in isolated yeast mitochondria by triggering recombination-mediated replication. Nucleic Acids Res. 37, 749–761 (2009).

76. C. M. Grant, F. H. MacIver, I. W. Dawes, Mitochondrial function is required for resistance to oxidative stress in the yeast Saccharomyces cerevisiae. FEBS Lett. 410, 219–222 (1997).

77. A. Couce, O. A. Tenaillon, The rule of declining adaptability in microbial evolution experiments. Front. Genet. 6 (2015), doi:10.3389/fgene.2015.00099.

