## Supplementary Materials for "Genetically controlled mtDNA editing prevents ROS damage by arresting oxidative phosphorylation"

**This PDF file includes:**

Materials and Methods

Supplementary Text S1 to S6

Figs. S1 to S10

Table S1

Captions for Data S1 to S30

**Other Supplementary Materials for this manuscript include the following:**

Data S1 to S30 (<https://data.mendeley.com/datasets/mvx7t7rw2d/draft?a=95381e47-dc80-47af-85ab-e0478912a209>)

Materials and Methods

Yeast cells

*Founder strain:* We used a single, haploid clone of the *S. cerevisiae* strain YPS128 (*MAT****α*** *ura3*::*NatMX*-barcode *ho::HYGMX*) as the background genotype. YPS128 is a wild, oak isolate with a North American genome composition (*49*) whose respiratory capacity has not been impaired by domestication (*17*). In contrast to common lab-strains, it carries neither *HAP1* defects, impairing mitochondrial regulation, nor *MIP1* defects that lead to spontaneous mtDNA loss (*50*).

*Deletion strains:* We generated a *ρ*^0^ YPS128 strain, lacking all mtDNA, by deleting *MIP1*. We also constructed gene deletion strains lacking *SOD2*, *SOD1*, *CCP1*, *ATG32*, *YAP1*, *RTG2* and *RTG3*. Genotypes: YPS128, *MAT****α,*** *ura3*::*natMX*-barcode, *ho::hygMXΔ, genex::kanMX,* with each target deleted from start to stop codon*.* We also constructed a *mip1Δsod2Δ* double deletion mutant as *sod2::kanMX, mip1::URA3.* For all strains, 2-3 independent clones, verified by PCR to carry the deletion cassette and to lack the deleted gene at the target locus, were isolated and retained. Several independent clones were isolated for all constructs, and used as replicates in all experiments.

*Aneuploidic strains:* Cells with and without one extra chromosome II, III or V were generated by repeatedly (3x) backcrossing clones from endpoint (*t_50_*) populations carrying chromosome duplications to founder clones of the opposite mating type (*MAT****a*** *ho::HYGMX*). Each backcross was done on YPD (Yeast Peptone Dextrose) medium using haploids verified by qPCR to retain the chromosome duplication. Diploid hybrids were selected after three days of growth on solid minimum media (0.675% Yeast Nitrogen Base (CYN2210, ForMedium), 2% (w/v) D-Glucose, pH=6-6.5 (NaOH), 2.5% agar) medium and sporulated overnight on solid 1% potassium acetate sporulation medium to generate recombined haploids. These were genotyped at the *URA* and *ho* locus and *ura- MAT****α*** haploids were passed on to the next round of backcrossing. After three rounds of backcrossing, we selected *ura- MAT****α*** haploids with (*n=*2 clones) and without (*n=*2 clones) the chromosome duplication of interest and estimated their respective growth rates (*n=*6) in a completely randomized design on the media of interest. We compared the cell doubling time of clones with and without the respective chromosome duplication.

*Cox4-EGFP fusion strain:* To construct Cox4-EGFP fusion as a reporter for mitochondrial morphology, founder cells were transformed with PCR fragments of EGFP amplified from pYM27, with flanking regions homologous to *COX4.* Downstream of *COX4* we inserted kanMX as selection marker during transformation. Cox4 localizes to the mitochondrial inner membrane (*51*).

Cell cultivation media

Except where otherwise stated, yeast strains were cultivated on a Complete Supplement Mixture medium (CSM medium; hereafter: “Background medium”) composed of 0.14% Yeast Nitrogen Base (CYN2210, ForMedium), 0.50% NH_4_SO_4_, 0.077% Complete Supplement Mixture (CSM, DCS0019, ForMedium), 2.0% (w/w) glucose, pH set to 5.80 with 1.0% (w/v) succinic acid and 0.6% (w/v) NaOH. For solid medium cultivations, 2.0% (w/v) agar was added. For pre-cultures to glycine, isoleucine, citrulline and tryptophan selection environments, we modified the background medium to avoid confounding growth on the stored nitrogen (*52*) by replacing CSM by 20 mg/L uracil and by reducing the NH_4_SO_4_ concentration (30 mg N/L). Simple modifications of the background medium were made to generate four of the eight stressor environments: +0.8 µg/mL rapamycin, +400 µg/mL paraquat (methylviologen; N,N-dimethyl-4-4′-bipiridinium dichloride) (*18*), +3 mM arsenic ([As III]; NaAs_2_O_3_), +62.5 mg/L citric acid. To generate the four other stressor environments, we replaced NH_4_SO_4_ in the background medium with 30 mg N/L of L-glycine, L-isoleucine, L-citrulline or L-tryptophan, together with 20 mg/L uracil. For the respiratory growth experiments, we replaced 2% glucose with 2% glycerol. For the H_2_O_2_ growth experiments, we added +3mM H_2_O_2_ to the background medium. We cast all solid plates 10-15 h prior to use in PlusPlates (Singer Instruments, UK), on a level surface, by pouring 50 mL of selection medium in the same upper right corner of each plate. We removed excess liquid by drying plates in a laminar airflow in a sterile environment. We stored cells at -80 °C in 20% glycerol and cultivated them at 30°C. Populations were subsampled and transferred to and from plates using robotics (ROTOR HDA, Singer Instruments Ltd, UK), at the indicated transfer format.

Experimental evolution of cells

We single streaked and then expanded a single haploid YPS128 clone to moderate colony size (~2 million cells), sampled the colony (~50 000 cells) and expanded the sample until stationary phase (~2 million cells; 36h) in 5 mL of background medium. A subsample of these founder cells were stored. We poured a sample of the stationary phase culture on top of a solid plate (background medium) and allowed the lawn of cells to grow, again until stationary phase (72 h). We then repeatedly sampled the lawn using 384 short pin pads to generate eight solid plates with 1152 colonies each. These colonies served as pre-cultures (*t_-1_*) to the first selection cycle of each of the eight selection environments. We expanded these pre-cultures on background, or nitrogen background, medium until stationary phase (~2 million cells; 72 h). We transferred samples of the pre-culture with 384 short pin pads to experimental plates to generate the 1152 populations to be evolved in each selection environments (table S1). We then cycled all 8x1152 populations through 50 rounds of expansion until stationary phase (72 h), subsampling and transfer to fresh plates, to produce *t_1_* to *t_50_*. We evolved many (*n*=24-192, see figure legends) *mip1Δ, ccp1∆,* *yap1∆*, *sod1∆, sod2∆* and *atg32∆* cell populations in a similar design, over a varying number of growth cycles. We consistently interleaved several wild type cell populations to serve as controls on the same plates.

Establishing and cultivating frozen chronological records of cell populations

In parallel to the sampling of colonies for transfer to fresh plates, we systematically sampled a large subset of populations in each environment to generate a frozen chronological record of their evolution. For each of the eight selection environments, we systematically sampled (1536 short pin pads) the same 96 populations at the end of growth cycles 0, 1, 2, 3, 4, 5, 7, 9, 12, 15, 20, 25, 30, 35, 40, 45 and 50, to generate a dense chronological adaptation record of 768 populations. We transferred the samples to a liquid selection medium (100 μL), expanded the populations until stationary phase (72 h), added 100 μL of glycerol (final concentration: 20% (w/w)) and stored them at -80 °C. We thawed and re-suspended these frozen stocks, and transferred cells (96 short pin pads) to a solid background, or a nitrogen background medium. To generate a randomized design, we used the *randint* function in the Python package NumPy (version 1.15.4). We pre-cultivated cells until stationary phase (72 h), sampled and transferred pre-cultures (1536 short pin pads) to selection environments plates, interleaving (384 short pin pads) 384 separately pre-cultivated, wild type, founder controls among the evolving populations on each plate. We cultivated all 1536 cells populations until stationary phase (72 h) while tracking their growth and adaption as described below. Using the same design, we also established and cultivated a frozen chronological record of paraquat-exposed *mip1Δ, ccp1∆,* *yap1∆*, *sod1∆, sod2∆* and *atg32∆* cell populations.

We performed three distinct release-from-selection experiments, using the frozen chronological records as start point. First, we thawed, re-suspended, sampled and transferred *t_0_*, *t_1_*, *t_2_*, *t_3_*, *t_4_*, *t_5_*, *t_7_* and *t_50_* samples of the 96 frozen paraquat-adapting populations to no stress solid medium plates. We evolved these populations over ten (~84 generations) growth cycles on no stress plates, sampled each population at the end of each growth cycle and stored samples at -80 °C (as above) to create a chronological record with samples first adapted to paraquat for different time-periods, and then released from the paraquat selection, again for different time periods. We thawed, re-suspended, sampled, randomized and pre-cultivated (no stress) this second chronological record, and sampled and transferred stationary phase cells to paraquat selection plates. Second, to compare the kinetics of loss of paraquat adaptive gains to that of populations adapting to other challenges characterized by fast adaptation, we repeated (3x) the above selection relaxation experiment, but including not only the 96 stored paraquat adapting populations but also those adapting to arsenic and glycine. We selected the time point in the chronological record where the populations had achieved 70-90% of their endpoint adaptation. We then thawed, re-suspended and sampled these stocks, expanded revived cells under relaxed selection for 10 growth cycles and created a frozen chronological record, which was revived, randomized, pre-cultivated and cultivated in the original stress, as above (*n=*5). For the glycine-adapting populations a nitrogen-limited background medium was used. Third, to compare the kinetics of loss of paraquat adaptation to that of the restoration of both intact mtDNA and of respiratory growth, we again repeated the release-from-paraquat experiment, but only for the five sequenced paraquat-adapting populations (A7, A8, B12, B5 and B8). Procedures were as above, but we replicated the experiment for each sample x3 and assayed both paraquat and respiratory (2% glycerol) growth at *n=*5 (randomization) for each replicate.

Tracking cell growth and adaptation

*Counting cells in growing populations:* We assayed the growth of cell populations in all experiments using the Scan-o-matic system (*53*), version 1.5.7 (https://github.com/Scan-o-Matic/scanomatic.git). Cultivation plates were maintained undisturbed and without lids for the duration of the experiment (72 h) in high-quality desktop scanners (Epson Perfection V800 PHOTO scanners, Epson Corporation, UK) standing inside dark, temperature (30.0 °C) and moisture controlled thermostatic cabinets with air circulation. We imaged plates at 20 min intervals using transmissive scanning at 600 dpi, identified the position of colonies and extracted intensities for pixels included in, and outside, each colony. For each colony, we estimated its sum pixel intensity as well as the median pixel intensity of the local background, subtracted the latter from the former and converted the remaining cell-associated pixel intensity to cell counts by using a pre-established calibration function, which had been obtained by estimating cell numbers using both spectrometry and flow cytometry. We smoothed and quality controlled growth curves, rejecting approximately 0.3% of growth curves as erroneous while being blinded to sample identities (for details, see Zackrisson et al. (*53*)). To allow direct visual comparison of growth curves of different samples while accounting for confounding effects from initial population size differences, we adjusted growth curves shown in figures in the *y*-dimension. We applied the function
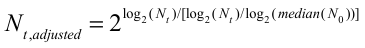
 to the mean growth curves to be visualized in figures, where *N_0_* is the mean initial population size across replicates, *N_t_* is the mean population size at time *t* across replicates, and the median(*N_0_*) is the median of the mean *N_0_* of the samples to be visualized together.

*Cell doubling time and adaptation:* We extracted the cell doubling time, *D,* from expanding cell populations. We used the 384 fixed spatial controls introduced at every fourth position to account for systematic doubling time variations within and across plates. By interpolating across the log_2_(*D*) values of the 384 measured controls (see (*53*), we estimated the log_2_(*D*) value a control colony would have had in each position. From the log_2_(*D*) value for each colony we then subtracted the corresponding log_2_(*D*) control value, thereby obtaining a normalized, relative log_2_ doubling time, log_2_(*D*)*_norm_*. Log_2_(*D*)*_norm_* is reported in Supplemental Information Data S2, S3, S5, S6, S7, S8, S23, S24, S29 and S30. When relevant, we also adjusted the log_2_(*D*)*_norm_* value for the bias associated with spatial controls having a slightly different pre-cultivation history than evolving populations, by use of the equation log_2_(*D*)_adj_ = log_2_(*D_t_* )*_norm_* - log_2_(*D_0_* )*_norm_*, where the subscripts *t* and *0* refer to the growth cycle number. Log_2_(*D*)_adj_ is reported in Supplemental Information Data S1. In some cases, we converted *D_norm_* back to a doubling time in hours while maintaining the normalization in order to ease interpretation. This measure is denoted *Doubling* *time* in figures, and set equal to *2^Dnorm^D_control, grand_*, where *D_control, grand_* is the grand mean of the raw doubling times of all controls run in a particular experimental series.

*Counting cell generations:* We estimated the number of cell generations for any missing growth cycle by interpolating the values estimated for the two adjacent growth cycles. For each cell population the total number of cell generations was calculated by summing over all growth cycles.

*Maximum possible reduction in cell doubling time:* We estimated how much of the maximum possible reduction in cell doubling time the paraquat adapting populations had achieved at a given generation number by comparing their cell doubling times with that of the founder population growing on normal medium, assuming that the latter represented a lower boundary for what was physiologically possible.

RNA sequencing to measure *SOD1*, *SOD2* and *CCP1* expression

Wild type cell populations were pre-cultivated for two consecutive 72 h growth cycles on no stress background medium, sampled and transferred to background medium w. and w/o 400 µg/mL paraquat (as above). We exposed cells to paraquat for three growth cycles and then removed the paraquat for one additional growth cycle. We sampled cell populations: i) immediately (10-15 s) after transfer (paraquat cycle 1 and 2), after 0.75 h (paraquat cycle 1 and 2), 1.5h (paraquat cycle 1), 5 h (paraquat cycle 1 and 2), 20 h (paraquat cycle 1) and 25 h (paraquat cycle 1). Cells to be harvested at early time-points (<5 h after transfer) were cultivated in a 6144 colony format, otherwise we used a 1536 format. All samples corresponding to the same growth cycle were cultivated in parallel. To generate one replicate of one sample, we harvested all colonies on a plate by pouring 5mL of liquid medium, w. or w/o paraquat, on top of the solid medium and scrapping off colonies with a sterile plastic rake into this liquid medium. The cells were pelleted at 12000 G (2 min in 4 °C), re-suspended in RNAlater (Sigma Aldrich R0901) and stored at 4 °C. We extracted RNA from all the stored samples in parallel, first diluting the RNAlater solution with an equal volume of PBS and then pelleting cells at 5000G (5 min, 4 °C). Cells were lysed by adding 600 µL of acid washed 0.5 mm beads and subsequent homogenization in a FastPrep homogenizer (three rounds a 40 s at 6 m s^-1^ separated by 1 min on ice). RNA quality was determined using a Tapestation 2200 and Nanodrop (threshold; ABS_260/280_ > 2.2 and RINe > 8). RNA sequencing was performed at SciLife (Stockholm, Sweden) using the Illumina TruSeq Stranded mRNA kit and a NovaSeq 6000 S4. RNA reads were checked for contamination using FastQ Screen (*54*). Filtered reads were aligned to the YPS128 reference genome using STAR (*55*), and optical duplicates were marked with Picard-tools. The abundance of the *SOD1*, *SOD2* and *CCP1* transcripts was quantified with featureCounts from the subread package across all samples (*56*). We normalized their read counts as fragments per kilobases per million reads, using the DESeq2 package for R (*57*). We estimated significant differences compared to no stress at *t*_0_ using Wald tests and Benjamini-Hochberg FDR correction, with a cut-off of *q*<0.05. The normalized read counts for *SOD1*, *SOD2* and *CCP1* are reported in Data S19.

DNA sequencing of evolving cell populations

*Long read (PacBio) sequencing of the YPS128 founder strain:* The total genomic DNA was extracted from a founder population cultivated overnight in background medium, using a standard phenol-chloroform protocol. We sequenced the genome on a PacBio RS II instrument using the P4-C2 chemistry. Additional PacBio sequencing data of the same YPS128 genotype were incorporated from and older assembly (*58*). A total of 9 SMRT cells were used to produce 1352628 reads, corresponding to approximately 205x genome coverage. We ran the *de novo* assembly using the hierarchical assembly protocol RS_HGAP_Assembly3.3 with an expected genome size of 12 Mb. Data were deposited at Sequencing Read Archive (SRA), accession number PRJNA622836.

*Very long read (Oxford nanopore) sequencing*: To exclude confounding effects of very early mtDNA changes, i.e. during freezing, thawing and the first round of paraquat cultivation, we thawed and subsampled frozen lawn cells (A7 lawn position) and cultivated these in the presence of paraquat until stationary phase. DNA was extracted using Qiagen Genomic-tip 100/G DNA extraction kit. Libraries for Oxford Nanopore sequencing were prepared using 1D Native barcoding genomic DNA with the EXP-NBD104 and SQK-LSK108kit. The flow cell version was FLO-MIN106, and the raw nanopore reads were basecalled by guppy (v2.1.3) with a minimal quality score cutoff of 5 (options: --qscore_filtering --min_qscore 5). For all basecalled reads that passed the quality filter, demultiplexing was further performed by guppy with the help of the guppy_reads_classifier.pl from LRSDAY (v1.3.1)(*59*). The de-multiplexed reads were processed by LRSDAY (v1.3.1) for adapter trimming, reads down sampling (down sampled to 50X coverage), *de novo* assembly, assembly polishing, assembly scaffolding, and dotplot visualisation. We deposited data at Sequencing Read Archive (SRA), accession number PRJNA622836.

*Resequencing of adapted populations and populations released from selection*: We thawed and subsampled frozen chronological record populations and cultivated cells in liquid medium in presence of paraquat overnight (24 h). DNA was extracted using a modified protocol of the Epicentre MasterPure Yeast DNA Purification Kit. Pool sequencing was performed at SciLife (Stockholm, Sweden), using Illumina HiSeq2500, 2x126 bp. Libraries were prepared using the Nextera XT kit to accommodate the low DNA yield from small cultures. At least two founder controls were included in each flow cell.

*Calling de novo point mutations:* Sequenced reads were quality-trimmed and nextera transposase sequences were removed with TrimGalore (v.0.3.8). Reads were mapped to the YPS128 pacbio assembly (see above) using BWA MEM (v.0.7.7-r441). PCR and optical duplicates were flagged using Picard-tools (v.1.109 [1716]). Base alignment quality scores were calculated using samtools calmd (v.0.1.18 [r982:295]) and variants were called using Freebayes (v0.9.14-8-g1618f7e). All alleles were reported regardless of frequency or genotype model. Variants were annotated using SnpEFF (v.3.6c). Variants below a quality score of 20 and variants present in the sequenced founder samples were filtered out. Data were deposited at Sequencing Read Archive (SRA), accession number PRJNA622836, and reported in Data S9.

*Calling aneuploidies:* Aneuploidies were called using a sliding, non-overlapping 200bp window coverage of reads mapped. Reads with a MAPQ of <1 were not counted. The window coverage ratio was calculated as
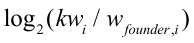
 where
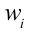
is the depth of coverage of mapped reads in each 200bp window, $w_{founder}$ is the depth of coverage of each *i* in a founder sequenced in the same flow cell and
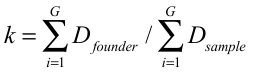
, where *G* is the YPS128 genome size and *D* is the depth of coverage for each nucleotide. Aneuploidies were called by determining the median log_2_ window coverage for each chromosome. Data are reported in Data S10 and S11.

*Calling mtDNA copy number change:* mtDNA copy number was calculated for each sample using a sliding, non-overlapping window of 1 kB. The mtDNA copy number relative to the euploid nuclear genome was calculated for each window as:
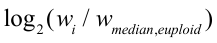
, where is the median of all 1 kB windows of the nuclear genome, excluding chromosomes with detected aneuploidies. We estimated the median absolute number of mtDNA molecules across all windows, assuming one copy of the nuclear genome and no sequencing bias for mitochondrial DNA, as
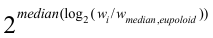
. These are reported in Data S12, S13 and S14.

Numerical model of evolving cell populations

*Cell population parameters:* To generate simulated adaptation trajectories based on empirical effect sizes and mutation rates of point mutations and aneuploidies, we used an individual-based model implemented in Python (*21*). We repeated each simulation 1152x. We started from a haploid, isogenic founder population that was subsampled at the end of each growth cycle to found the next cultivation cycle. The population parameters were population size at the start of each growth cycle (*N*), the number of cell divisions before subsampling in each growth cycle (*M_t_*) and the total number of growth cycles (*n=*50 cycles). When the total population size reached 2*^Mt^N* cells, *N* cells were sub-sampled randomly to found the next cycle. *N* was set to equal the approximate mean across all empirical sub-samplings. *M_t_* was set to equal the mean (across populations) empirical measure in each growth cycle *t.* Each cell divided 12x before it died. Mating, meiosis, sporulation or ploidy change were not included, and there was no population structure.

*Mutation effect sizes:* We estimated the mutation effect sizes empirically. To estimate gene loss-of-function mutation effect sizes underlying simulations shown in Figure 1D, we used the haploid BY4741 single gene deletion collection (*MAT***a**;*his3Δ1;leu2Δ0;met15Δ0;ura3Δ0; genex::kanMX*) (*60*), which was cultivated in absence and presence of each stressor, wherefrom the cell doubling times were extracted. Collection size: *n=*4580, each cultivated at *n=*6. We report the doubling time data for this collection in Data S7. To estimate chromosome duplication effect sizes, we used the duplications of chromosome II, III, V, X and XVI constructed by backcrossing (as above). We reconstructed the chromosome duplications IV, VI, VIII, IX, XI, XII, XIII, XIV and XV as in (*61*). We genetically modified the founder clone genotype to match the *his3Δ* (complete deletion by transformation with pSH47) and *can1::STE2pr-HIS3* genotype of the aneuploidic construct, as described in Zebrowski *et al*., 2008 (*61*), and used these as controls. Duplications of I and VII could not be obtained by either methods, despite repeated tries. Strains carrying duplications were cultivated in absence and presence of each stressor (*n=*9) and doubling times, *D*, extracted. We report these data in Data S8.

*Mutation rate parameters:* All cells began as identical, haploid founder cells. Cells had 4947 nuclear encoded protein genes, and 16 chromosomes specified by the sequenced reference genome (R64-1-1). Cells had no mitochondrial genome. Essential genes were not included. Cells independently and randomly acquired nuclear genome mutations as chromosome duplications and point mutations in protein coding genes at the end of each cell division. Mutation rates were constant, equal for all genomes, for all chromosomes and for all nucleotide sites. Chromosomes and nucleotide sites were only allowed to mutate once. Sites on new chromosomes did not mutate. Chromosome duplications occurred at rate of *µ*=4.85*10^-5^ duplications/cell division. Point mutations occurred at a rate *µ*=0.33*10^−9^ point mutations/bp/division (*62*).

*Mutation effect size parameters:* We tracked the mutations of each cell, its reproductive age, and its cell division time. Mutations and cell division time were passed to daughter cells. Cells began at a cell division time equal to the founder population doubling time. Change in cell division time was affected by mutations only, and mutations only affected cell division time. Because chromosome duplication and loss-of-gene function point mutations are the most common drivers of experimental adaptation (*63*), we assumed these to be the only sources of change in cell division time. We estimated the cell division effect size of chromosome duplications as described above. We estimated the cell division effect size of point mutations by downloading the SIFT yeast database (http://sift-db.bii.a-star. edu.sg/public/Saccharomyces_cerevisiae/EF4.74/) and extracting all possible stop gain base changes and nonsynonymous mutations, with attached SIFT scores (*64*). All stop gain base changes and all nonsynonymous mutations with a SIFT score <0.05 affected cell division time with an effect size equal to the population doubling time effect of the corresponding gene deletion Reproductive age did not affect cell division time and there were no cell-cell interactions. We assumed that the cell doubling time under stress at any instance could not become shorter than the measured mean founder cell doubling time in absence of stress. We implemented the well documented principle of diminishing return of mutations with increasing fitness (*65*, *66*), by letting a mutation *m* define the cell division time, *D_m_*, obtained from the equation
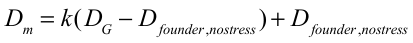
, where
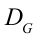
 is the cell division time of the genotype before the mutation occurred, and
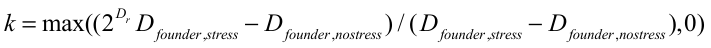
.
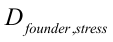
 is the measured mean doubling time of the founder in presence of stress, and *D_r_* js the estimated doubling time effect size of the mutation assuming no epistasis. No other form of epistasis was included.

Quantitative PCR of mtDNA genes in evolving cell populations

To track the copy number dynamics of mtDNA genes in evolving populations, we performed quantitative PCR (qPCR) on the frozen chronological samples from a subset of populations (A4, A7, A8, A9, B5, B8, B12, D1). The samples were revived in 3 mL liquid background media supplemented with 400 µg/mL paraquat, expanded to stationary phase (72 h). DNA was extracted from harvested cells using a MasterPure Yeast DNA purification kit (Epicentre), as per the manufacturer’s instructions. Primers were designed for each of the protein and rRNA encoding mtDNA genes and for one nuclear control: *CDC5*. The small size and extreme AT richness of the RNA subunit of RNase P (*RPM1*) prevented the design of working PCR primers for this mtDNA gene. We ran qPCR for duplicates of the entire frozen chronological record of one mtDNA gene in a single run, together with *CDC5* controls. The qPCR was performed using iTaq Universal SYBR Green Supermix (total volume: 20 µL) and run on a Bio-Rad CFX Connect using Bio-Rad Hard-Shell PCR 96-well thin-wall plates sealed with adhesive transparent film. The PCR protocol was: initial denaturation (95 °C; 15 min) followed by 45 cycles of: denaturation (95 °C; 15 s), anneal (60 °C; 30 s), extension (72°C; 30 s), and a melting curve analysis. We quantified PCR products at the annealing step of each cycle due to the low melting temperature of all PCR products, which follows from extremely low GC% of the mtDNA. The relative copy number was calculated as
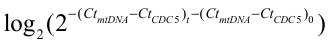
.

We capped all Ct values at 30. We also set Ct values to 30 for rare (5.6%) sample replicates where the *COX1* primer pair produced non-PCR based background signals. We report the data in Data S15 and S29. Primers used are reported in Data S18.

Light and fluorescence microscopy of evolving cells

We performed light and fluorescence microscopy on DNA (DAPI) stained $\rho^{+}$ (founder, WT), $\rho^{-}$ (population A7 at *t_50_*), $\rho^{--}$ (population B8 at *t_50_*) and $\rho^{0}$ (*mip1Δ;* lacking the mitochondrial DNA polymerase) cells to validate that $\rho^{--}$ cells contained mitochondrial DNA. We cultivated cell populations overnight in liquid background medium, diluted pre-cultures in fresh media to OD_600_=0.3 and incubated with agitation in 15 mL growth tubes until OD_600_=0.6. Cells were pelleted by centrifugation at 3000 G for 1.5 min. The supernatant was discarded and cells were suspended in 1 mL of ice cold 70% ethanol. Cells were fixated by incubation in RT for 5 min. The cells were then pelleted again by centrifugation, washed in non-ionic water and re-suspended in PBS (Fisher Bioreagents BP2944-100) solution. DNA was stained with DAPI (D3571, Merck) at 50 ng/mL just before imaging. Images were acquired using a Zeiss Axio Observer Z1 Inverted microscope with Plan-Apochromat 100x/1.40 Oil DIC M27 objective and AxioCam MR R3 camera.

Electron microscopy of evolving cells

We assayed mitochondrial dynamics by electron microscopy before $O_{2}^{-}$ stress, after 5 h (~1 cell doubling)of paraquat stress, during long-term paraquat stress (A7, *t_50_*) and 5 h after release from long-term paraquat stress (A7, *t_50_*). Frozen stocks were revived by transfer to solid background medium (with or without paraquat) and cells were cultivated for 5 h (1 population doubling). Cells were harvested by rinsing the plate with 5 mL liquid background media (with or without paraquat), suspended by stirring with a plastic spreader and pelleted by centrifugation at 600 G for 1.5 min. Pelleted cells were frozen under high pressure using a Wohlwend Compact 03 (M. Wohlwend GmbH, Sennwald, Switzerland). Freeze substitution was performed in a Leica EM AFS2 (Leica Microsystems, Vienna, Austria) by incubating the cells with 2% uranyl acetate dissolved in 10% methanol and 90% acetone for 1 h at -90 °C (*67*). Freeze substituted cells were washed (2x) in 100% acetone and the temperature was raised 2.9 °C/h to -50 °C. Cell pellets were broken into smaller pieces to improve resin infiltration. Infiltration of cells was performed using a ladder of Lowicryl HM20 (Polysciences, Warrington, PA) diluted in decreasing acetone concentrations (1:4, 2:3, 1:1, 4:1) followed by three changes in pure Lowicryl. Each step lasted 2 h. The resin was polymerized with UV light, first for 72 h at -50 °C and then for 24 h at room temperature. Resin embedded blocks were sectioned in 70 nm ultra-thin sections using a Reichert-Jung Ultracut E Ultramicrotome (C. Reichert, Vienna, Austria) equipped with an ultra 45° diamond knife (Diatome, Biel, Switzerland). Sections were collected on copper grids coated with 1% formvar and stained with 2% uranyl acetate and Reynold’s lead citrate (*68*). Stained sections were imaged at 120 kV using a Tecnai T12 microscope (FEI Co., Eindhoven, The Netherlands) and a Ceta CMOS16 camera. The IMOD package (*69*) was used for quantification of 100 cell sections per sample. To validate quantifications, a large subset of images was analyzed by blind-test by a second person. Conflicting quantifications were discarded. We report these data in Data S16 and S17.

Confocal microscopy of evolving cells

We also tracked the mitochondrial dynamics by confocal microscopy of fluorescently labelled (Cox4-EGFP) mitochondria in cells before exposure to paraquat, after 7 h (1 doubling) of paraquat stress, after 79 h (1 growth cycle + 1 doubling) of stress and 7 h after release from 144 h (2 growth cycles) of paraquat exposure. We isolated 3 transformants and ensured that their respiratory growth was normal by spot-assays on glycerol medium. Transformants were cultivated with or without paraquat for 0 h or 72 h in liquid medium, the stationary phase cultures were diluted in fresh media with or without paraquat to OD_600_=0.3 and then incubated with agitation until OD_600_=0.6 (exponential phase). The cells were then pelleted by centrifugation at 3000 G for 1.5 min and washed in MQ water, centrifuged and suspended in PBS. Cells were fixated by incubation at room temperature with 3.7% formaldehyde, washed 3x in PBS, suspended in ProLong Diamond mounting media and directly mounted on slides and imaged in the microscope. *Z*-stacks of cells were acquired using a Zeiss Axio Observer LSM 700 inverted confocal microscopy with a Plan-Apochromat 63x/1.40 Oil DIC M27. The signal was averaged between 4 frames to reduce noise. Z-stacks were color-coded according to Z-dimension (slice) using Temporal-Color Code in Fiji/ImageJ v. 1.52. We pre-processed images using the difference of Gaussians (DoG) to enhance the objects to be measured, i.e. either cells or mitochondria. To measure cells, we calculated the 3D gradient of the DoG filtered images, using triangle algorithm. To measure mitochondria, we calculated the 3D median filter of the DoG filtered images before segmentation. We segmented images using Otsu’s method, separating touching objects processed using seed-assisted watershed algorithm. The used seeds were suggested automatically, with a human blinded to sample identities correcting for mistakes. The final quantification of shape descriptors for each cell and the mitochondria within were done in MATLAB, with the relevant code and readme files available at: https://github.com/CamachoDejay/SStenberg_3Dyeast_tools. We report the data in Data S20.

**Resource availability:** Sequence data that support the findings of this study have been deposited in Sequencing Read Archive (SRA) with the accession codes PRJNA622836. The growth phenotyping code can be found at <https://github.com/Scan-o-Matic/scanomatic.git>, the simulation code at <https://github.com/HelstVadsom/GenomeAdaptation.git> and the imaging code at <https://github.com/CamachoDejay/SStenberg_3Dyeast_tools>. The authors declare that all other data supporting the findings of this study are available within the paper as Supplemental Information Data S1-S30, which can be previewed at https://data.mendeley.com/datasets/mvx7t7rw2d/draft?a=95381e47-dc80-47af-85ab-e0478912a209.

**Materials availability:** All unique strains and stored populations generated in this study are available from the Lead Contact without restriction.

Supplementary Text

Supplementary Text 1

The chosen paraquat dose caused the cell doubling time to increase 2.5-3 fold. Under this condition, copper/zinc dependent O_2_^•−^ dismutase (Cu/ZnSOD, Sod1), manganese dependent O_2_^•−^ dismutase (MnSOD, Sod2) and mitochondrial cytochrome C peroxidase (Ccp1) transcript levels increased 2-12 fold in early lag-phase, and the cells maintained the elevated expression of these antioxidant transcripts throughout the exponential growth phase and the following two growth cycles (fig. S1) without causing any reduction of the cell doubling time. Assuming that this substantially enhanced expression also reflect mobilization of the whole repertoire of primary antioxidant defenses, this demonstrates that the paraquat-induced O_2_^•−^ production was well above the reach of these defenses.

Supplementary Text 2

We measured the mtDNA content by short-read genome sequencing in five random populations exposed to paraquat. We found a clear inverse temporal association between the median mtDNA coverage and the reduction in cell doubling time (Figure 3A). However, because of the repetitive nature of the yeast mtDNA, the short-read data did not provide a reliable description of the temporal mitochondrial genome dynamics.

Supplementary Text 3

$\rho^{0}$ cells devoid of mtDNA show a 2-fold decrease in cellular O_2_^•−^ production also in supposedly fermenting yeast cells growing on glucose in an aerobic environment (*27*), suggesting that oxidative phosphorylation was far from being totally repressed when cells became exposed to paraquat (*70*).

Supplementary Text 4

Considering that the first line of defense against enhanced levels of intramitochondrial O_2_^•−^ is its dismutation into hydrogen peroxide (H_2_O_2_) by O_2_^•−^ dismutases, and that H_2_O_2_ is a major redox signaling agent through specific protein targets (*71*), the sensing and signaling mechanisms triggering mtDNA editing could involve H_2_O_2_. To investigate this possibility, we measured the effect of H_2_O_2_ on cell doubling time in 96 cell populations that had been exposed to paraquat for 50 growth cycles (*t_50_*). We found that they grew slower than founder cell populations exposed to the same concentration of H_2_O_2_ (fig. S6A), which implied that the paraquat adaptation had enhanced the sensitivity to H_2_O_2_. Moreover, cells lacking the cytochrome C peroxidase Ccp1, a key mitochondrial peroxidase (*72*), and cells lacking the transcription factor Yap1, which controls the transcriptional antioxidant response to H_2_O_2_ (*73*, *74*), adapted as wild type cells to paraquat (fig. S6B). Together with the observations that yeast cells react to H_2_O_2_ by amplifying their mtDNA (*75*) and that $\rho^{0}$ cells are H_2_O_2_-sensitive (*76*), these data suggest that H_2_O_2_ did not play a significant role during the adaptation to paraquat.

Supplementary Text 5

Cells lacking either one of the O_2_^•−^ dismutases showed virtually no growth when exposed to the original paraquat dose (400 μg/mL) (fig. S6C). As the adaptation kinetics depends heavily on the strength of selection (*77*), we reduced the paraquat concentration such that mutant cells were somewhat less or slightly more (12.5 μg/mL and 50μg/mL for *sod2Δ*) stressed than wild type cells at 400 μg/mL. We found that the *sod2Δ* populations mostly displayed no (12.5 μg/mL) or much slower (50 μg/mL) doubling time reduction over 10 growth cycles. The *sod1Δ* cells (12.5 μg/mL) adapted as wild type cells (Fig. 4A). Based on these results, we chose to use a paraquat concentration of 50 μg/mL to track the pattern of mtDNA deletions in eight *sod2Δ* cell populations by qPCR.

Supplementary Text 6

Nuclear point mutations rarely recurred in the same genes across these endpoint populations (fig. S9B). Moreover, in the five populations for which we had time-resolved sequencing, we found that the nuclear point mutations failed to coincide in time with the $O_{2}^{-}$ adaptation (fig. S9C). These nuclear point mutations were therefore unlikely to have played a prominent causative role in the adaptation process. However, all but four endpoint populations carried extra copies of chromosome II (*n=*29), III (*n=*21) and/or V (*n=*16) (fig. S10A) at near fixation (mean *p*: 0.97). These chromosome gains succeeded the early and very swift $O_{2}^{-}$ adaptation phase (fig. S10B). To assess their contribution to the second phase of adaptation, we generated clones carrying the individual aneuploidies in a wild type background (see Methods) and compared the tolerance for paraquat and the capacity for respiratory growth with and without extra chromosomes. We found that the duplications of chromosomes II and V reduced the cell doubling time during paraquat exposure by 31 and 38 min, respectively (fig. S10D). Duplication of chromosome III caused no apparent reduction in cell doubling time. Assuming an additive phenotypic effect of the chromosome II and V duplications, the cell doubling time would be reduced by 69 min. Together with the effect from the mtDNA deletions disrupting OXPHOS function, this would correspond to cells realizing 81% of the possible adaptation. As the cells actually realized on average 72.6%, an approximately additive phenotypic effect of these duplications appears to fully explain the second phase of adaptation in populations having both duplications genetically fixed. Moreover, the chromosome II and V duplications contributed marginally (~2.4 h and 1.6 h, respectively) to the complete loss of respiratory growth (fig. S10C).

Supplementary Figures

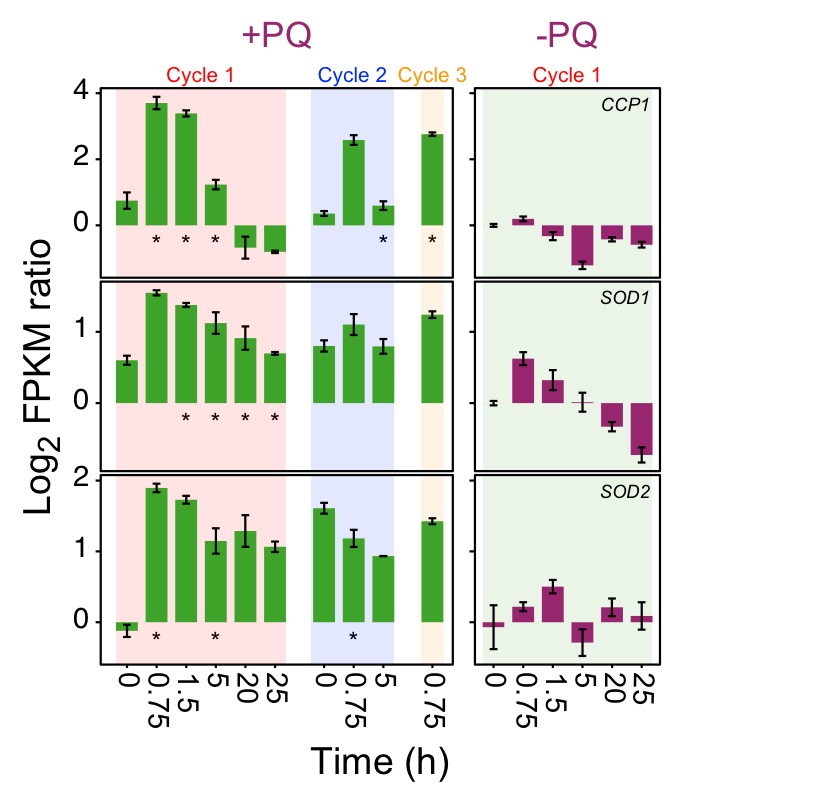

Fig. S1. Titration of the paraquat (PQ) dose to inundate the primary antioxidant defenses.

*Left panel:* Color columns show the mRNA expression (FPKM; Fragments Per Kilobase Million) of *CCP1*, *SOD1* and *SOD2,* during the first, second and third growth cycle in the presence of 400 μg/mL paraquat. Note that the cells have not yet been exposed to paraquat at time, *t*=0 in Cycle 1. *Right panel:* mRNA expression in the founder population in a paraquat-free growth medium. *x*-axis: time (h) in each growth cycle. *y*-axis: Expression values are shown as a log_2_ ratio in relation to expression in a paraquat-free medium at *t*=0. Error bars: S.E.M (*n=*3). * = significant (Wald test, FDR *q*=0.05) difference.

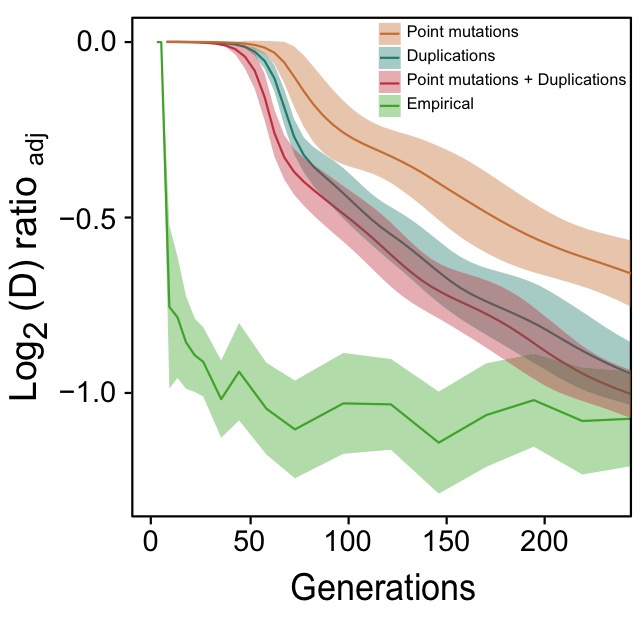

**Fig. S2 Comparison of predicted paraquat adaptation with the experimental data.** The experimental adaptation data (green) on 96 populations (each measured at *n*=6) are the same as in Fig. 1A. The three prediction graphs generated by the numerical model are each based on 1152 replicate runs. Shade: S.D.

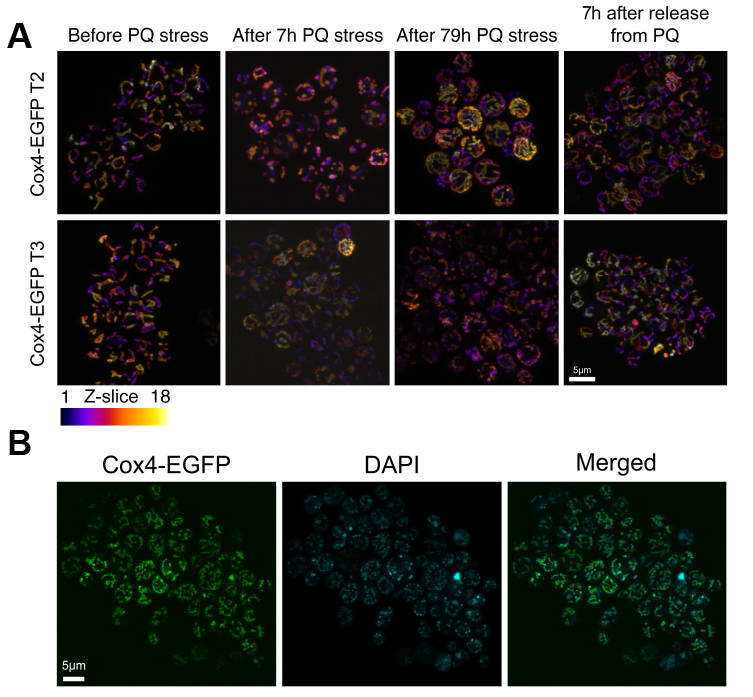

**Fig. S3. Paraquat (PQ) stress leads to rapid mitochondrial fragmentation.**

(**A**) Confocal microscopy of yeast cells with a Cox4-GFP inner mitochondrial membrane marker. Panels show cells before PQ exposure, after 7 h of exposure, after exposure over one growth cycle (72 h + 7 h), and 7 h after release from two growth cycles under PQ stress. Color: *z*-dimension (yellow=front, black=back), samples were sliced in 18 slices with the first slice being closest to the camera. Two Cox4-EGFP transformants, isolated independently from the one in Fig. 2B, are shown. (**B**) Confocal micrographs of cells exposed to PQ for 7 h, showing the predominating presence of mtDNA in the mitochondrial fragments.

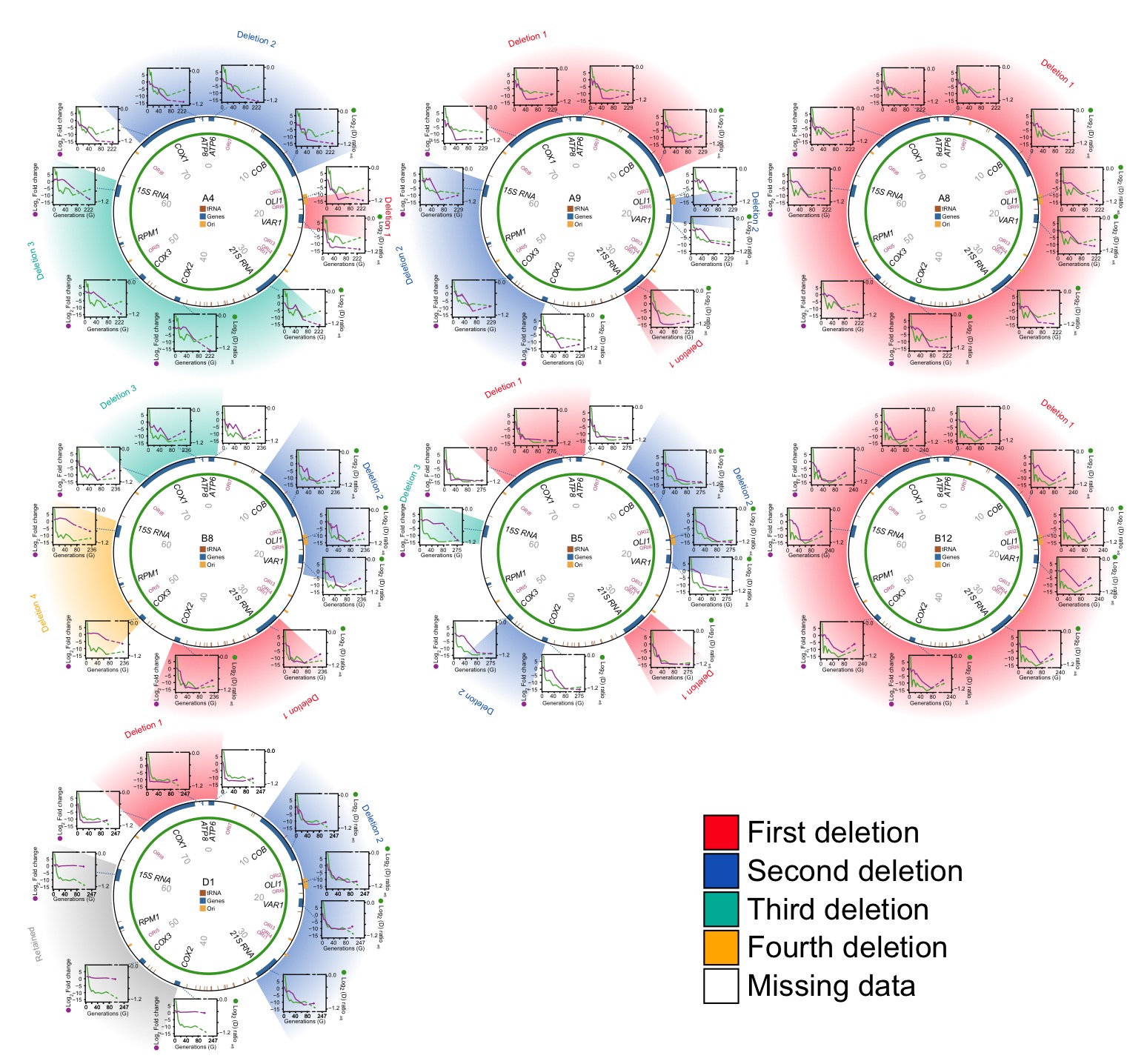
**Fig. S4. Editing of mtDNA during the early adaptation to paraquat.**

*Circle:* mtDNA (77 kB) before exposure to paraquat stress, with genes, known origins of replication and position (numbers) indicated. *Diagrams:* mtDNA copy number (left *y*-axis, qPCR, purple line) of each protein and rRNA encoding gene, and the associated temporal adaptation profile (right *y*-axis, adjusted log*_2_*(*D*) ratio relative to founder*,* green line), in populations A4, A9, A8, B8, B5, B12 and D1. Error bars: S.E.M. (*n=*2). Colored fields describe concomitant copy number.

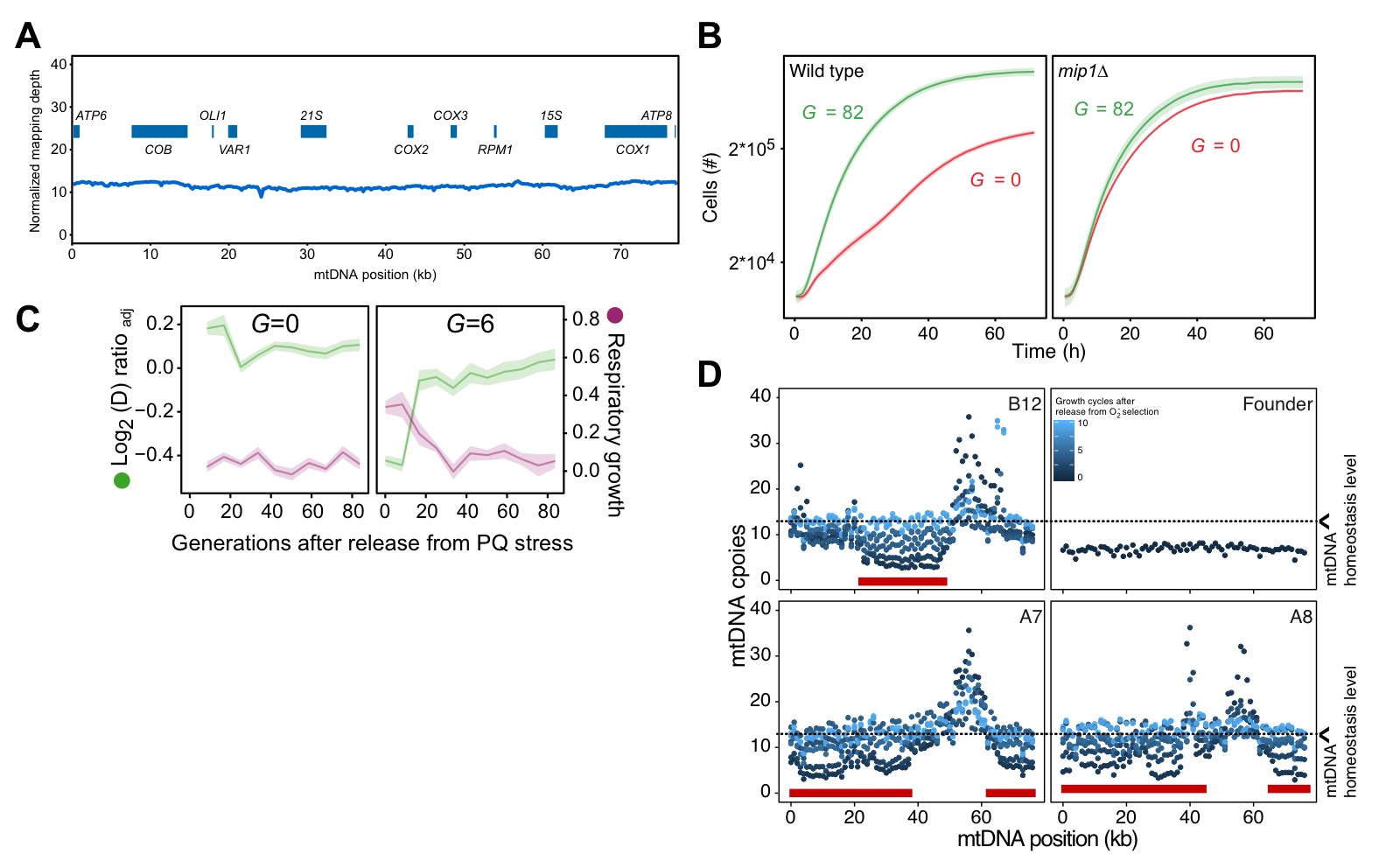

**Fig. S5. Homeostatic restoration of mtDNA copy numbers and ability for respiration after release from paraquat (PQ) stress.**

(**A**) mtDNA copy numbers at pre-adaptation levels. *y*-axis, median mtDNA coverage, in 0.5kb windows, relative to that of the haploid nuclear genome. *x*-axis: nucleotide position. Data is from long read sequencing. (**B**) Comparisons of the growth of wild type (mean of *n=*64 populations; left panel) and *mip1Δ* (mean of *n=*24 populations; right panel) cells on paraquat (400 μg/mL), before (*G* =0) and after 10 growth cycles (mean*=*82 generations) of adaptation to paraquat. Shade: S.E.M across cell populations (each measured at *n*=3). (**C**) Respiratory (glycerol) growth (right *y*-axis, purple line, log_2_ doubling time relative to founder), and loss of adaptation (left *y*-axis, green line), in cell populations not exposed to paraquat (G=0) (left panel, founder) and cell populations exposed to 6 generations of paraquat exposure (right panel) and then released from this selection over *G* generations (x-axis) of growth in absence of paraquat. The mean of five cell populations (A7, A8, B5, B8 and B12; same as in Fig. 3A), each measured at *n=*5, is shown. Shade: S.E.M. (**D**) Recovery of mtDNA copy numbers (*y*-axis, median coverage in 1 kb windows relative to the euploid nuclear genome) in cell populations released from 6 generations of paraquat exposure (dot color: growth cycles after release from paraquat). Red bars: positions of mtDNA deletions.

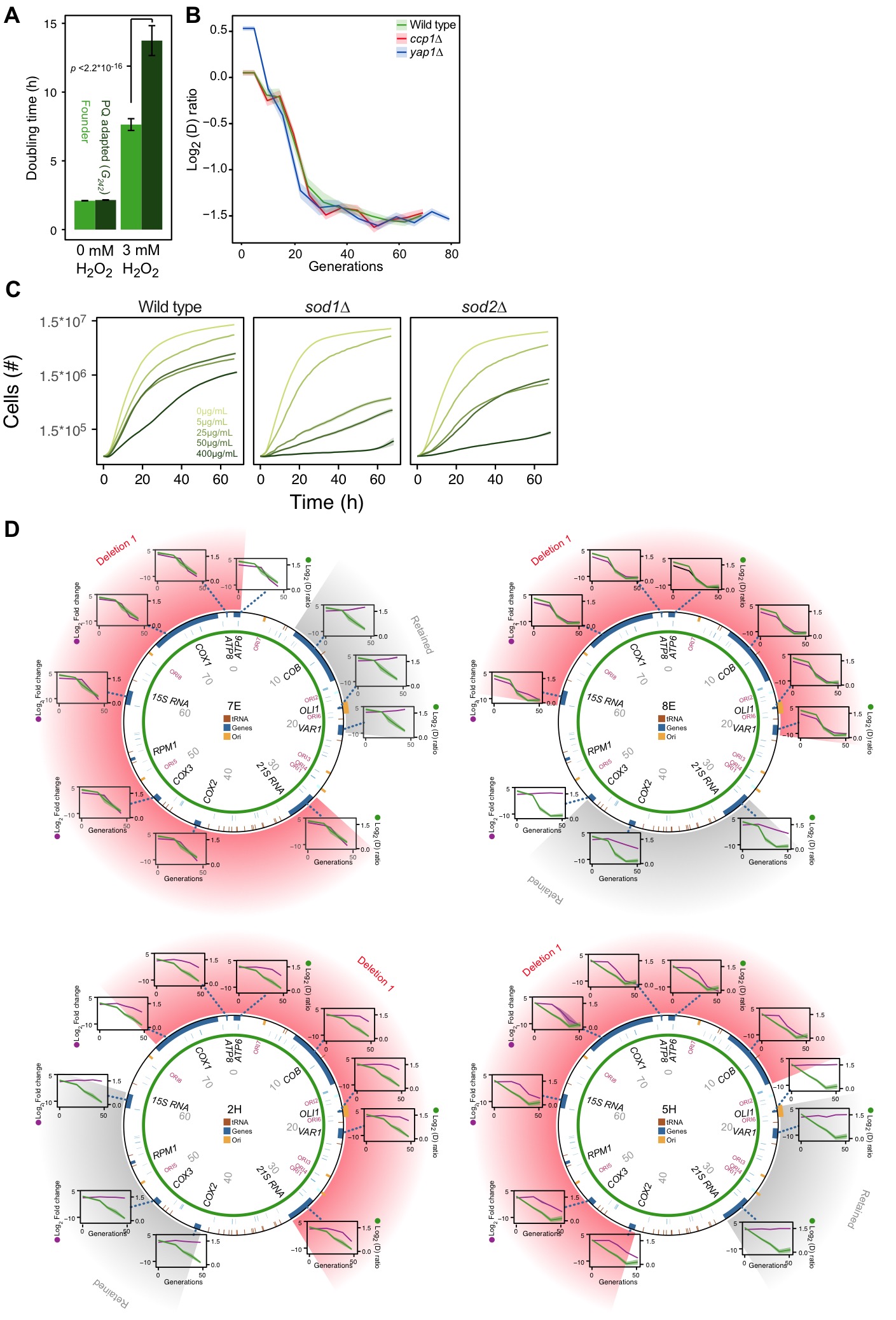

**Fig. S6. The mtDNA deletion process critically involves Sod2, but not Sod1 or H_2_O_2_**

(**A**) Cell doubling time (h) of founder (light green) and paraquat adapted (*G*=242 (mean), dark green) cell populations, in the presence of 0 mM and 3 mM H_2_O_2._ Error bars: S.E.M. (*n=*768 cell populations). *p*-values: Welch two-sided t-test. (**B**) Adaptation profiles of *ccp1Δ* and *yap1Δ* cell populations (*n*=12-16 populations, each measured at *n=*3) exposed to 400 μg/mL of paraquat. Shade: S.E.M. (**C**) Comparison of the growth of wild type (left), *sod1Δ* (center) and *sod2Δ* (right) cell populations in increasing (color intensity) doses of paraquat. Shade: S.E.M. (*n=*96). (**D**) mtDNA deletions in four *sod2Δ* cell populations adapting to 12.5 μg/mL of paraquat. *Circle:* mtDNA (77 kb) before paraquat exposure. Genes, origins of replication and nucleotide positions (kb) are indicated. Colored fields describe mtDNA deletions with concomitant copy number change. *Diagrams:* mtDNA copy number change (left *y*-axis, purple line, (*n=*2) of individual mtDNA genes during adaptation to paraquat (right *y*-axis, green line) in *sod2Δ cell* populations 7E, 8E, 2H and 5H. Shade: S.E.M.

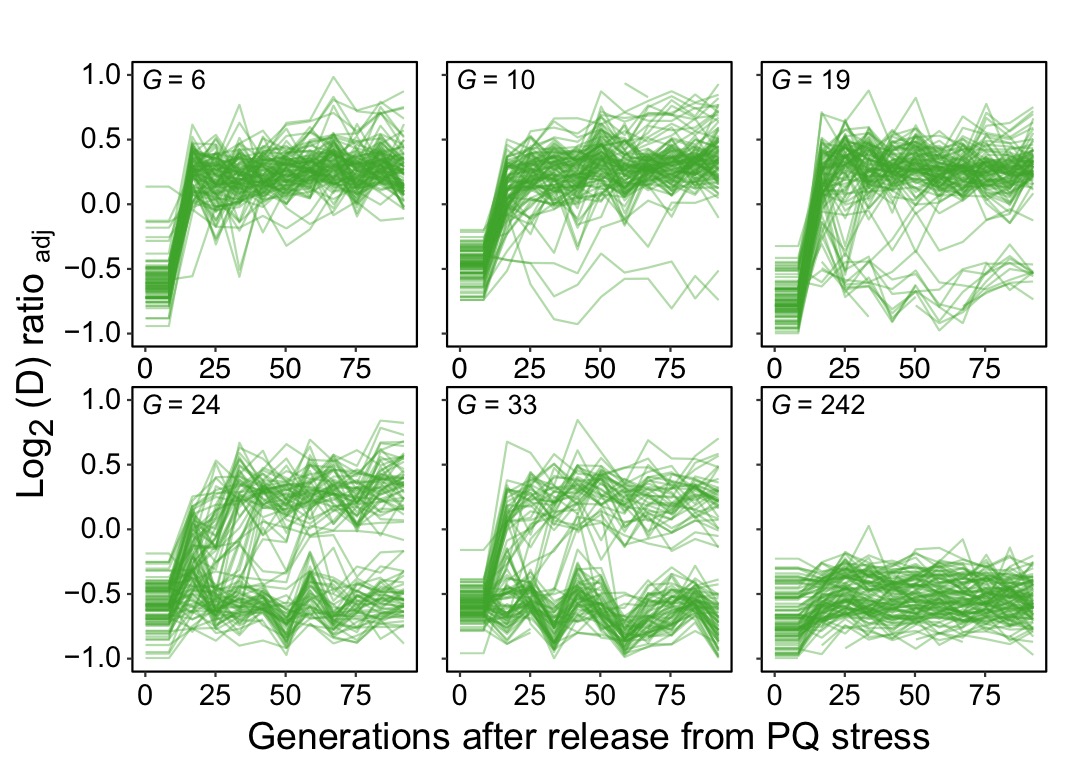

**Fig. S7. Long-term exposure to paraquat causes genetic fixation.**

We released 96 populations from paraquat exposure after 6, 10, 19, 24, 33 and 242 generations (means) of adaptation (*panels*), and by re-exposing the populations to paraquat after a given number of generations of growth on a paraquat-free medium we could monitor the fraction of populations where the paraquat adaptation had become genetically fixed. Lines: 96 populations (each measured at *n*=5). Note that after 6 generations of adaptation, all populations rapidly lose their acquired paraquat adaptation, implying no genetic fixation, while after 242 generations the adaptation has become genetically fixed in all populations.

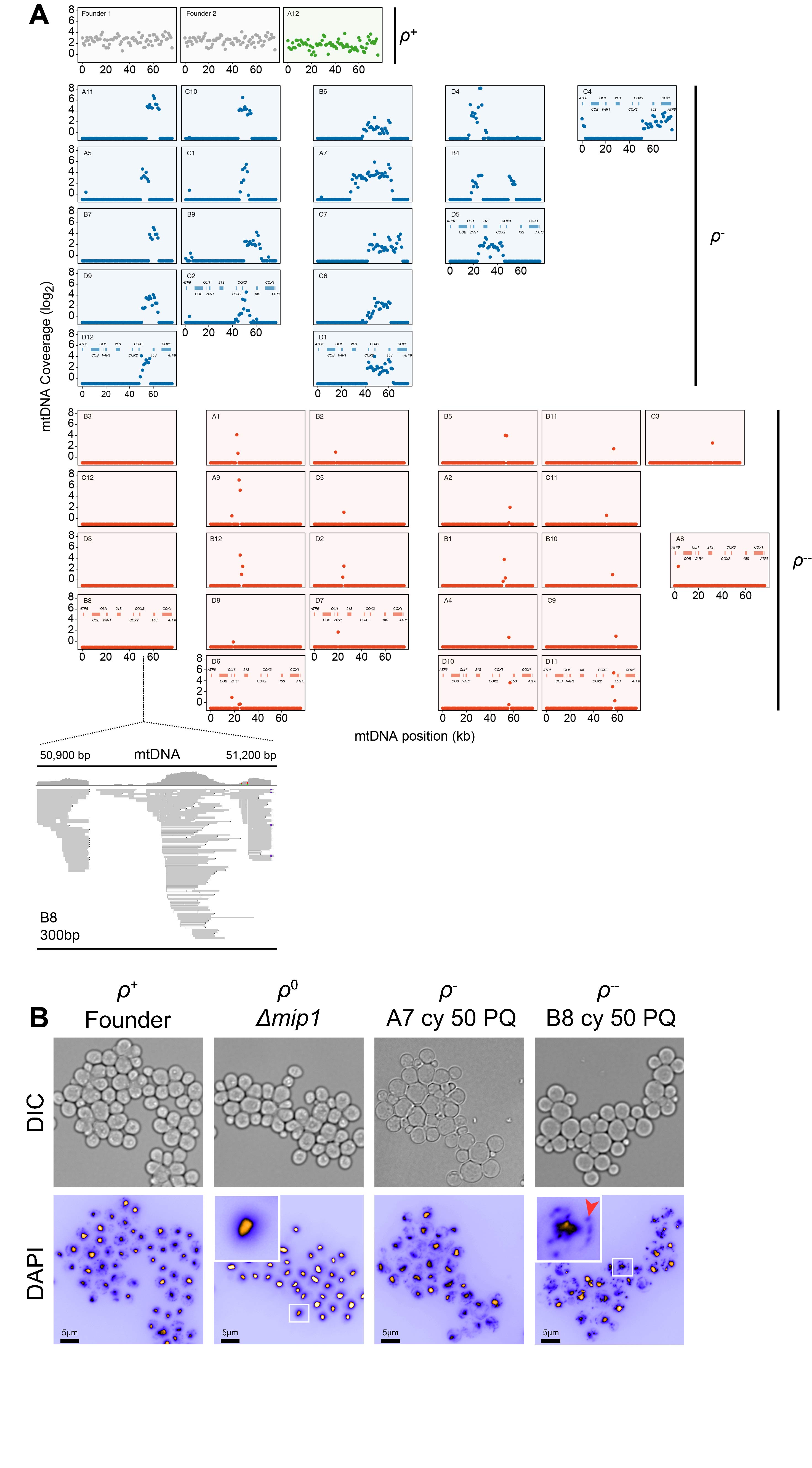

**Fig. S8.** **mtDNA loss during long-term exposure to paraquat.**

(**A**) Sequenced populations (*n=*44) adapted (*t_50_*) to chronic paraquat exposure were classified (color) as $\rho^{+}$ (mtDNA intact, green, *n=*1),
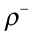
 (6-30 kb mtDNA segments retained, blue, *n=18*) and
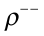
 (<2 kb mtDNA segments retained, red, *n=25*). *y*-axis: mtDNA copy number (median coverage in 0.5kb windows relative to the euploid nuclear genome). *x*-axis: mtDNA position. *Below*: gene positions. *Read map:* a zoom-in on a 300 bp mtDNA stretch which is mapped to by the
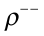
 population B8 mtDNA sequence reads. (**B)**
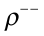
 cells with low mtDNA sequence coverage retain very short (<300 bp) mtDNA segments. *Micrographs:* Light (top; DIC) and fluorescence (bottom) microscopy of DAPI stained DNA in
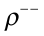
 (B8 at *t_50_*),
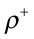
 (founder),
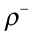
 (A7 at *t_50_*), and true
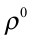
 (*mip1Δ*) cells. *Insets:* Zoom-in on indicated
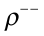
and true
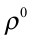
cells. Note that
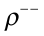
 cells contain mitochondrial DNA (red arrow) while true
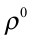
cells do not.

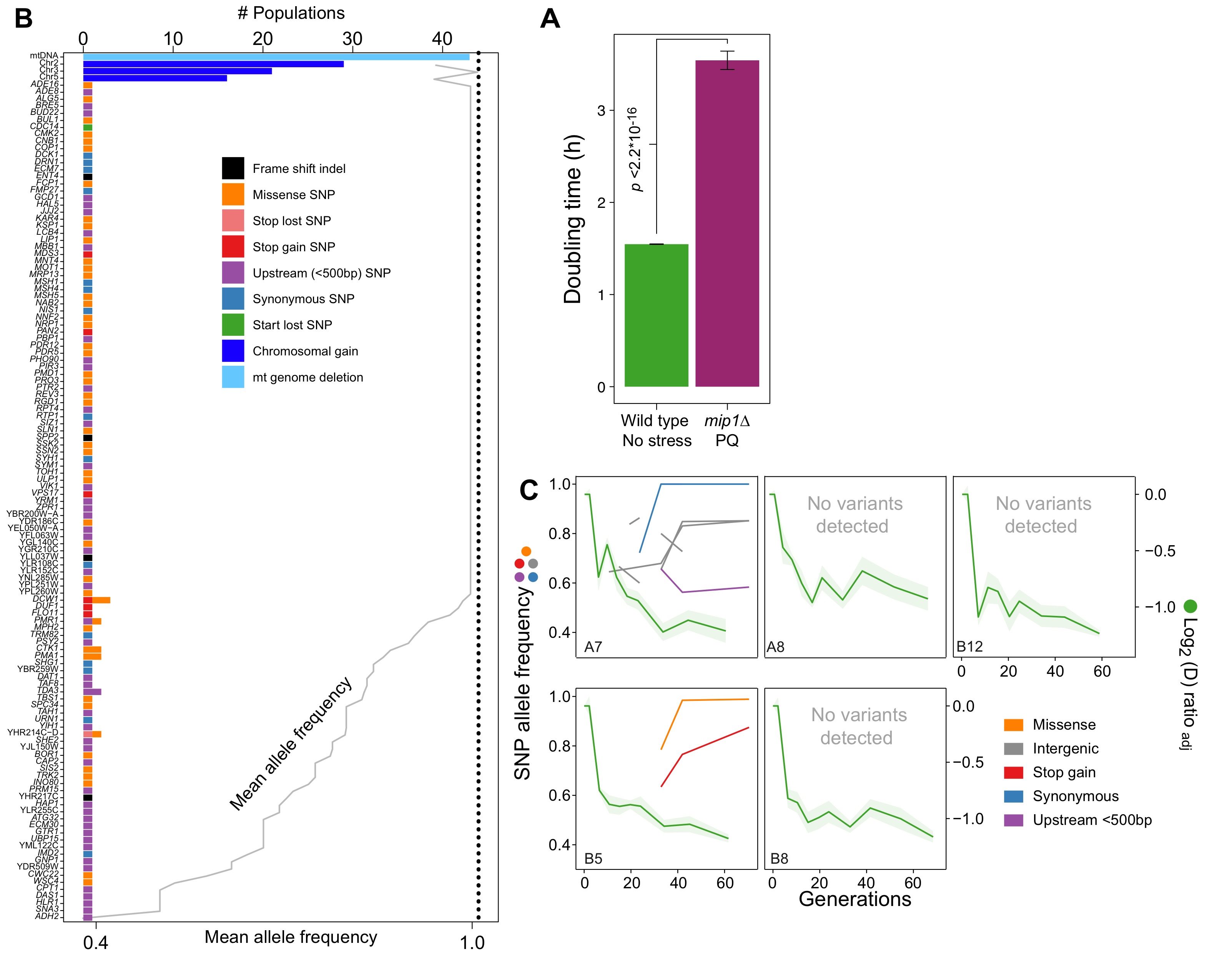

**Fig. S9.** **Nuclear genome evolution during long-term exposure to paraquat (PQ).**

(**A**) Doubling time (h) of *mip1Δ* cell populations (*n=*432) growing in the presence of 400 μg/mL of paraquat, compared to that of founder cell populations (*n=*768) growing on ordinary medium. *p*-values Welch two-sided t-test. (**B**) mtDNA deletion and chromosome II, III and V duplications recur across populations adapted to long-term paraquat stress, but nuclear genes with point mutations rarely do. Upper *x*-axis: Number of populations in which a gene carries *de novo* point mutations, a chromosome is duplicated or a mtDNA segment is deleted. Dotted line: number of sequenced populations. Lower *x*-axis (grey line): For genes containing SNPs, the line shows the mean allele frequencies of SNPs in the gene. For chromosome or mtDNA copy number variations the line shows the mean sequence coverage across the chromosome or mtDNA relative to that of the haploid nuclear genome. *y*-axis: Genes with point mutations, chromosomes with duplications and mtDNA. Bar color: Type of variation. (**C**) The early, swift adaptation to paraquat (right *y*-axis, green line, *A,* shade: S.E.M., *n=*6) precedes point mutations (left *y*-axis, non-green lines, allele frequency). *x*-axis: Generations of exposure to paraquat. *Panels:* Sequenced populations (A7, A8, B5, B8, and B12; same as in Fig. 3A). Line color: variant type. Variants pre-dating adaptation, supported by few (<10) reads or (<2) time points or shared across environments (>2) were filtered out.

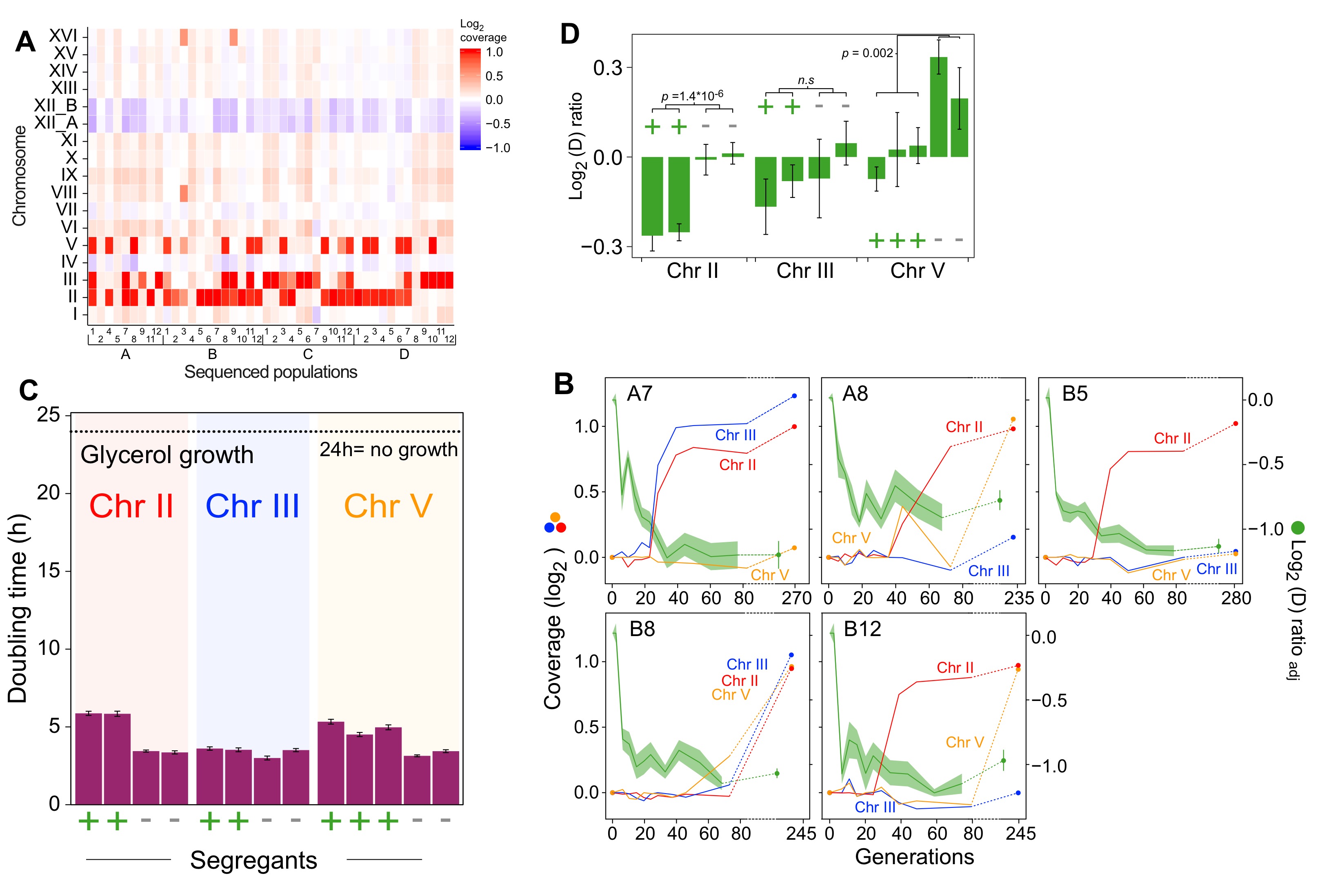

**Fig. S10. Chromosome duplications explain the second phase of adaptation to paraquat.**

(**A**) Chromosome II, III and V duplications are common after 50 cycles (mean of *G*=242 generations) of paraquat exposure. Color: Chromosome copy number (log_2_ median coverage relative to haploid nuclear genome. (**B**) Chromosome II, III and V duplications appear in the second phase of paraquat adaptation. *Panels:* five populations (A7, A8, B5, B8, and B12). Left *y*-axis (non-green lines): Chromosome copy number (log_2_ of median sequence coverage across the chromosome relative to the median of the nuclear genome). Color: chromosome (II=blue, III=red and V=yellow). Right *y*-axis (green line): paraquat adaptation, *A*. Shade: S.E.M. (*n=*6). Broken lines: no data. (**C-D**) Chromosome II and III duplications reduce the cell doubling time under paraquat stress **(C)** but cannot explain the complete loss of respiratory growth in the parent populations (**D**). We backcrossed (x3) cells adapted to 242 generations (*t_50_*) of paraquat exposure to founder cells over consecutive meioses and compared the growth on paraquat of 2-3 segregants with and without duplicated chromosome. *x*-axis: cells w. (+) and w/o (-) individual chromosome duplications. Error bars: S.E.M. (*n=*6). *p*-values: Welch two-sided t-test. Broken line: No growth, corresponding to doubling time >24 h (the measurement limit).

**Supplementary Tables**

**Table S1:** List of Selection Agents Used in Evolution Experiments

| **Selection** | **Compound** | **Concentration** | **Type** |
| --- | --- | --- | --- |
| Mitochondrial $O_{2}^{-}$ stress | Paraquat | 400 µg/mL | Mitochondrial $O_{2}^{-}$ producer |
| Arsenic (III) stress | Arsenite (NaAsO_2_) | 3 mM | Toxic metalloid |
| Rapamycin stress | Sirolimus | 0.8 µg/mL | TOR-inhibitor |
| Citric Acid stress | Citric Acid | 62.5 mg/mL | Weak organic acid |
| Glycine use | L-Glycine | 160.86 mg/mL (30 mg N/mL) | Nitrogen source |
| Citrulline use | L-Citrulline | 125.14 mg/mL (30 mg N/mL) | Nitrogen source |
| Tryptophan use | L-Tryptophan | 218.82 mg/mL (30 mg N/mL) | Nitrogen source |
| Isoleucine use | L-Isoleucine | 281.1 mg/mL (30 mg N/mL) | Nitrogen source |

**Captions for supplementary data S1 to S30**

**Data S1**

Doubling time data of 96 populations adapted to each of eight different environments over *G* generations; doubling times are bin the respective selection environment.

**Data S2**

Doubling time data of 96 populations adapted to paraquat for *G* generations exposed; doubling times are in paraquat and respiratory media (glycerol).

**Data S3**

Doubling time data of 96 populations adapted to paraquat for *G_s_* generations, followed by release from this selection over *G_r_* generations; doubling times on paraquat.

**Data S4**

Difference in doubling time in absence of stress and in the respective selection environment, for adapted populations having achieved 70-90% of their final adaptation.

**Data S5**

Doubling time data of 96 populations adapted to paraquat, arsenic and glycine over *G_s_* generations and then released from selection for *G_r_* generations; doubling times are in paraquat, arsenic and glycine respectively.

**Data S6**

Doubling time data of 96 populations adapted to paraquat for *G_s_* generations and then released from selection for *G_r_* generations; doubling times are in paraquat and respiratory media (glycerol).

**Data S7**

Doubling time data for the BY4741 single gene deletion collection under paraquat exposure; used as input for simulations in figure S2

**Data S8**

Doubling time data of disomic strains growing in paraquat; used as input for simulations in figure S2.

D**ata S9**

Small indels and SNPs called in sequenced paraquat adapted endpoint populations.

**Data S10**

Mean log_2_ coverage for each chromosome in each sequenced paraquat adapted endpoint population.

**Data S11**

Mean log_2_ coverage for each chromosome in five sequenced paraquat adapting populations over generations *G* of selection.

**Data S12**

Mean log_2_ coverage of 1 kb windows spanning the mitochondrial genome of each sequenced paraquat adapted endpoint population.

**Data S13**

Mean log_2_ coverage of 1 kb windows spanning the mitochondrial genome of five sequenced paraquat adapting populations over generations *G* of selection.

**Data S14**

Mean log_2_ coverage of 1 kb windows spanning the mitochondrial genome of sequenced populations adapting to paraquat and then released from this selection; data is given as a function of generations *G* of relaxation of selection.

**Data S15**

qPCR data for mitochondrial DNA genes and nuclear DNA controls over generations of paraquat adaptation

**Data S16**

Mitochondrial and cell area quantified, based on electron microscopy micrographs.

**Data S17**

Number of mitochondria quantified, based on electron microscopy micrographs.

**Data S18**

Primers used for strain construction and qPCR.

**Data S19**

FPKM data of selected oxidative defense genes, obtained from RNA-sequencing of cells exposed to paraquat.

**Data S20**

Number and volume of mitochondria of cells segmented from confocal microscopy micrographs of cells exposed to paraquat.

**Data S21**

Doubling time data of *mip1∆* cells grown in stress, and of wild type cells grown in no stress.

**Data S22**

Doubling time data of 96 endpoint populations adapted to paraquat; doubling times are in 0 and 3 mM of H_2_O_2_.

**Data S23**

Doubling time data of wild type and *atg32∆* populations adapting to paraquat over generations *G*; doubling times are in paraquat.

**Data S24**

Doubling time data of wild type, *rtg2∆, rtg3∆* and *mip1∆* populations adapting to paraquat; doubling times are in paraquat.

**Data S25**

Growth curves of wild type, *rtg2∆, rtg3∆* and *mip1∆,* adapted and not adapted to paraquat; doubling times are in respiratory media (glycerol).

**Data S26**

Doubling time data of wild type, *sod2∆ and sod1∆* populations adapting to paraquat; doubling times are in paraquat.

**Data S27**

Growth curves of populations of *mip1∆sod2∆, mip1∆, sod2∆* and wild type exposed to paraquat.

**Data S28**

Mean growth curves of wild type, *sod1∆* and *sod2∆* and wild type exposed to different concentrations of paraquat.

**Data S29**

qPCR data for mitochondrial DNA genes and nuclear DNA controls in *sod2∆* populations over generations of paraquat adaptation.

**Data S30**

Doubling time data of wild type, *ccp1∆ and yap1∆* populations adapting to paraquat; doubling times are in paraquat.

**References**

1.	N. Sun, R. J. Youle, T. Finkel, The Mitochondrial Basis of Aging. *Mol. Cell*. **61**, 654–666 (2016).

2.	H. Hu, C.-C. Tan, L. Tan, J.-T. Yu, A Mitocentric View of Alzheimer’s Disease. *Mol. Neurobiol.* **54**, 6046–6060 (2017).

3.	N. Ammal Kaidery, B. Thomas, Current perspective of mitochondrial biology in Parkinson’s disease. *Neurochem. Int.* **117**, 91–113 (2018).

4.	R. T. Hepple, Impact of aging on mitochondrial function in cardiac and skeletal muscle. *Free Radic. Biol. Med.* **98**, 177–186 (2016).

5.	J. M. T. Hyttinen, J. Viiri, K. Kaarniranta, J. Błasiak, Mitochondrial quality control in AMD: does mitophagy play a pivotal role? *Cell. Mol. Life Sci.* **75**, 2991–3008 (2018).

6.	K. J. Krishnan, A. K. Reeve, D. C. Samuels, P. F. Chinnery, J. K. Blackwood, R. W. Taylor, S. Wanrooij, J. N. Spelbrink, R. N. Lightowlers, D. M. Turnbull, What causes mitochondrial DNA deletions in human cells? *Nat. Genet.* **40**, 275–279 (2008).

7.	N. Nissanka, M. Minczuk, C. T. Moraes, Mechanisms of Mitochondrial DNA Deletion Formation. *Trends Genet.* **35**, 235–244 (2019).

8.	G. A. Fontana, H. L. Gahlon, Mechanisms of replication and repair in mitochondrial DNA deletion formation. *Nucleic Acids Res.* **48**, 11244–11258 (2020).

9.	J. J. Lemasters, Selective Mitochondrial Autophagy, or Mitophagy, as a Targeted Defense Against Oxidative Stress, Mitochondrial Dysfunction, and Aging. *Rejuvenation Res.* **8**, 3–5 (2005).

10.	A. S. Bess, T. L. Crocker, I. T. Ryde, J. N. Meyer, Mitochondrial dynamics and autophagy aid in removal of persistent mitochondrial DNA damage in Caenorhabditis elegans. *Nucleic Acids Res.* **40**, 7916–7931 (2012).

11.	K. Palikaras, N. Tavernarakis, Mitochondrial homeostasis: the interplay between mitophagy and mitochondrial biogenesis. *Exp. Gerontol.* **56**, 182–8 (2014).

12.	L. Sedlackova, V. I. Korolchuk, Mitochondrial quality control as a key determinant of cell survival. *Biochim. Biophys. Acta - Mol. Cell Res.* **1866**, 575–587 (2019).

13.	Å. B. Gustafsson, G. W. Dorn, Evolving and expanding the roles of mitophagy as a homeostatic and pathogenic process. *Physiol. Rev.* **99**, 853–892 (2019).

14.	H. Sies, C. Berndt, D. P. Jones, Oxidative Stress. *Annu. Rev. Biochem.* **86**, 715–748 (2017).

15.	T. Shpilka, C. M. Haynes, The mitochondrial UPR: Mechanisms, physiological functions and implications in ageing. *Nat. Rev. Mol. Cell Biol.* **19**, 109–120 (2018).

16.	M. Y. W. Ng, T. Wai, A. Simonsen, Quality control of the mitochondrion. *Dev. Cell*. **56**, 881–905 (2021).

17.	M. De Chiara, B. Barré, K. Persson, A. O. Chioma, A. Irizar, J. Schacherer, J. Warringer, G. Liti, *bioRxiv*, in press, doi:10.1101/2020.02.08.939314.

18.	H. M. Cochemé, M. P. Murphy, Complex I is the major site of mitochondrial superoxide production by paraquat. *J. Biol. Chem.* **283**, 1786–1798 (2008).

19.	P. R. Castello, D. A. Drechsel, M. Patel, Mitochondria are a major source of paraquat-induced reactive oxygen species production in the brain. *J. Biol. Chem.* **282**, 14186–93 (2007).

20.	X. Zou, B. A. Ratti, J. G. O’Brien, S. O. Lautenschlager, D. R. Gius, M. G. Bonini, Y. Zhu, Manganese superoxide dismutase (SOD2): is there a center in the universe of mitochondrial redox signaling? *J. Bioenerg. Biomembr.* **49**, 325–333 (2017).

21.	A. B. Gjuvsland, E. Zörgö, J. K. Samy, S. Stenberg, I. H. Demirsoy, F. Roque, E. Maciaszczyk-Dziubinska, M. Migocka, E. Alonso-Perez, M. Zackrisson, R. Wysocki, M. J. Tamás, I. Jonassen, S. W. Omholt, J. Warringer, Disentangling genetic and epigenetic determinants of ultrafast adaptation. *Mol. Syst. Biol.* **12**, 892 (2016).

22.	H. G. Sprenger, T. Langer, The Good and the Bad of Mitochondrial Breakups. *Trends Cell Biol.* **29**, 888–900 (2019).

23.	M. Frank, S. Duvezin-Caubet, S. Koob, A. Occhipinti, R. Jagasia, A. Petcherski, M. O. Ruonala, M. Priault, B. Salin, A. S. Reichert, Mitophagy is triggered by mild oxidative stress in a mitochondrial fission dependent manner. *Biochim. Biophys. Acta - Mol. Cell Res.* **1823**, 2297–2310 (2012).

24.	C. H.-L. Hung, S. S.-Y. Cheng, Y.-T. Cheung, S. Wuwongse, N. Q. Zhang, Y.-S. Ho, S. M.-Y. Lee, R. C.-C. Chang, A reciprocal relationship between reactive oxygen species and mitochondrial dynamics in neurodegeneration. *Redox Biol.* **14**, 7–19 (2018).

25.	Y. Liu, K. Okamoto, Regulatory mechanisms of mitophagy in yeast. *Biochim. Biophys. Acta - Gen. Subj.* **1865**, 129858 (2021).

26.	T. Lodi, C. Dallabona, C. Nolli, P. Goffrini, C. Donnini, E. Baruffini, DNA polymerase Î^3^ and disease: what we have learned from yeast. *Front. Genet.* **6** (2015), doi:10.3389/fgene.2015.00106.

27.	A. R. Reddi, V. C. Culotta, SOD1 integrates signals from oxygen and glucose to repress respiration. *Cell*. **152**, 224–35 (2013).

28.	Merriam-Webster, Edit. *Merriam-Webster’s Coll. Thes.*, (available at https://unabridged.merriam-webster.com/thesaurus/edit).

29.	C. B. Epstein, J. A. Waddle, W. Hale IV, V. Davé, J. Thornton, T. L. Macatee, H. R. Garner, R. A. Butow, Genome-wide responses to mitochondrial dysfunction. *Mol. Biol. Cell*. **12**, 297–308 (2001).

30.	T. Sekito, J. Thornton, R. A. Butow, Mitochondria-to-Nuclear Signaling Is Regulated by the Subcellular Localization of the Transcription Factors Rtg1p and Rtg3p. *Mol. Biol. Cell*. **11**, 2103–2115 (2000).

31.	B. A. Rothermel, A. W. Shyjan, J. L. Etheredge, R. A. Butow, Transactivation by Rtg1p, a Basic Helix-Loop-Helix Protein That Functions in Communication between Mitochondria and the Nucleus in Yeast. *J. Biol. Chem.* **270**, 29476–29482 (1995).

32.	B. A. Rothermel, J. L. Thornton, R. A. Butow, Rtg3p, a Basic Helix-Loop-Helix/Leucine Zipper Protein that Functions in Mitochondrial-induced Changes in Gene Expression, Contains Independent Activation Domains. *J. Biol. Chem.* **272**, 19801–19807 (1997).

33.	N. Guaragnella, L. P. Coyne, X. J. Chen, S. Giannattasio, Mitochondria–cytosol–nucleus crosstalk: learning from Saccharomyces cerevisiae. *FEMS Yeast Res.* **18** (2018), doi:10.1093/femsyr/foy088.

34.	G. Twig, A. Elorza, A. J. A. Molina, H. Mohamed, J. D. Wikstrom, G. Walzer, L. Stiles, S. E. Haigh, S. Katz, G. Las, J. Alroy, M. Wu, B. F. Py, J. Yuan, J. T. Deeney, B. E. Corkey, O. S. Shirihai, Fission and selective fusion govern mitochondrial segregation and elimination by autophagy. *EMBO J.* **27**, 433–446 (2008).

35.	T. Ban, T. Ishihara, H. Kohno, S. Saita, A. Ichimura, K. Maenaka, T. Oka, K. Mihara, N. Ishihara, Molecular basis of selective mitochondrial fusion by heterotypic action between OPA1 and cardiolipin. *Nat. Cell Biol.* **19**, 856–863 (2017).

36.	A. Kowald, T. B. L. Kirkwood, Transcription could be the key to the selection advantage of mitochondrial deletion mutants in aging. *Proc. Natl. Acad. Sci.* **111**, 2972–2977 (2014).

37.	D. Zorov, I. Vorobjev, V. Popkov, V. Babenko, L. Zorova, I. Pevzner, D. Silachev, S. Zorov, N. Andrianova, E. Plotnikov, Lessons from the Discovery of Mitochondrial Fragmentation (Fission): A Review and Update. *Cells*. **8**, 175 (2019).

38.	L. Aerts, V. A. Morais, in *Parkinson’s Disease* (Elsevier, 2017; https://linkinghub.elsevier.com/retrieve/pii/B978012803783600002X), pp. 41–75.

39.	S. Sabnam, H. Rizwan, S. Pal, A. Pal, CEES-induced ROS accumulation regulates mitochondrial complications and inflammatory response in keratinocytes. *Chem. Biol. Interact.* **321**, 109031 (2020).

40.	G. S. Nido, C. Dölle, I. Flønes, H. A. Tuppen, G. Alves, O. B. Tysnes, K. Haugarvoll, C. Tzoulis, Ultradeep mapping of neuronal mitochondrial deletions in Parkinson’s disease. *Neurobiol. Aging*. **63**, 120–127 (2018).

41.	B. Kalyanaraman, J. Joseph, S. Kalivendi, S. Wang, E. Konorev, S. Kotamraju, Doxorubicin-induced apoptosis: Implications in cardiotoxicity. *Mol. Cell. Biochem.* **234**–**235**, 119–124 (2002).

42.	G. Genc, A. Okuyucu, B. C. Meydan, O. Yavuz, O. Nisbet, M. Hokelek, A. Bedir, O. Ozkaya, Effect of free creatine therapy on cisplatin-induced renal damage. *Ren. Fail.* **36**, 1108–1113 (2014).

43.	S. S. Malhi, A. Budhiraja, S. Arora, K. R. Chaudhari, K. Nepali, R. Kumar, H. Sohi, R. S. R. Murthy, Intracellular delivery of redox cycler-doxorubicin to the mitochondria of cancer cell by folate receptor targeted mitocancerotropic liposomes. *Int. J. Pharm.* **432**, 63–74 (2012).

44.	Z. Song, H. Chang, N. Han, Z. Liu, Y. Liu, H. Wang, J. Shao, Z. Wang, H. Gao, J. Yin, He-Wei granules (HWKL) combat cisplatin-induced nephrotoxicity and myelosuppression in rats by inhibiting oxidative stress, inflammatory cytokines and apoptosis. *RSC Adv.* **7**, 19794–19807 (2017).

45.	M. Songbo, H. Lang, C. Xinyong, X. Bin, Z. Ping, S. Liang, Oxidative stress injury in doxorubicin-induced cardiotoxicity. *Toxicol. Lett.* **307**, 41–48 (2019).

46.	J. F. Moruno-Manchon, N. Uzor, S. R. Kesler, J. S. Wefel, D. M. Townley, A. S. Nagaraja, S. Pradeep, L. S. Mangala, A. K. Sood, A. S. Tsvetkov, Peroxisomes contribute to oxidative stress in neurons during doxorubicin-based chemotherapy. *Mol. Cell. Neurosci.* **86**, 65–71 (2018).

47.	X. Ren, J. T. R. Keeney, S. Miriyala, T. Noel, D. K. Powell, L. Chaiswing, S. Bondada, D. K. St. Clair, D. A. Butterfield, The triangle of death of neurons: Oxidative damage, mitochondrial dysfunction, and loss of choline-containing biomolecules in brains of mice treated with doxorubicin. Advanced insights into mechanisms of chemotherapy induced cognitive impairment (“chemobr. *Free Radic. Biol. Med.* **134**, 1–8 (2019).

48.	K. Adachi, Y. Fujiura, F. Mayumi, A. Nozuhara, Y. Sugiu, T. Sakanashi, T. Hidaka, H. Toshima, A Deletion of Mitochondrial DNA in Murine Doxorubicin-Induced Cardiotoxicity. *Biochem. Biophys. Res. Commun.* **195**, 945–951 (1993).

49.	G. Liti, D. M. Carter, A. M. Moses, J. Warringer, L. Parts, S. A. James, R. P. Davey, I. N. Roberts, A. Burt, V. Koufopanou, I. J. Tsai, C. M. Bergman, D. Bensasson, M. J. T. O’Kelly, A. van Oudenaarden, D. B. H. Barton, E. Bailes, A. N. Nguyen, M. Jones, M. A. Quail, I. Goodhead, S. Sims, F. Smith, A. Blomberg, R. Durbin, E. J. Louis, Population genomics of domestic and wild yeasts. *Nature*. **458**, 337–41 (2009).

50.	M. Gaisne, A. M. Bécam, J. Verdière, C. J. Herbert, A “natural” mutation in Saccharomyces cerevisiae strains derived from S288c affects the complex regulatory gene HAP1 (CYP1). *Curr. Genet.* **36**, 195–200 (1999).

51.	J. Zhu, Z.-T. Zhang, S.-W. Tang, B.-S. Zhao, H. Li, J.-Z. Song, D. Li, Z. Xie, A Validated Set of Fluorescent-Protein-Based Markers for Major Organelles in Yeast (Saccharomyces cerevisiae). *MBio*. **10**, 1–19 (2019).

52.	A. Gutiérrez, M. Sancho, G. Beltran, J. M. Guillamon, J. Warringer, Replenishment and mobilization of intracellular nitrogen pools decouples wine yeast nitrogen uptake from growth. *Appl. Microbiol. Biotechnol.* **100**, 3255–3265 (2016).

53.	M. Zackrisson, J. Hallin, L.-G. Ottosson, P. Dahl, E. Fernandez-Parada, E. Ländström, L. Fernandez-Ricaud, P. Kaferle, A. Skyman, S. Stenberg, S. Omholt, U. Petrovič, J. Warringer, A. Blomberg, Scan-o-matic: High-Resolution Microbial Phenomics at a Massive Scale. *G3 (Bethesda).* **6**, 3003–14 (2016).

54.	S. W. Wingett, S. Andrews, FastQ Screen: A tool for multi-genome mapping and quality control. *F1000Research*. **7**, 1338 (2018).

55.	A. Dobin, C. A. Davis, F. Schlesinger, J. Drenkow, C. Zaleski, S. Jha, P. Batut, M. Chaisson, T. R. Gingeras, STAR: ultrafast universal RNA-seq aligner. *Bioinformatics*. **29**, 15–21 (2013).

56.	Y. Liao, G. K. Smyth, W. Shi, featureCounts: an efficient general purpose program for assigning sequence reads to genomic features. *Bioinformatics*. **30**, 923–930 (2014).

57.	M. I. Love, W. Huber, S. Anders, Moderated estimation of fold change and dispersion for RNA-seq data with DESeq2. *Genome Biol.* **15**, 550 (2014).

58.	J.-X. Yue, J. Li, L. Aigrain, J. Hallin, K. Persson, K. Oliver, A. Bergström, P. Coupland, J. Warringer, M. C. Lagomarsino, G. Fischer, R. Durbin, G. Liti, Contrasting evolutionary genome dynamics between domesticated and wild yeasts. *Nat. Genet.* **49**, 913–924 (2017).

59.	J.-X. Yue, G. Liti, Long-read sequencing data analysis for yeasts. *Nat. Protoc.* **13**, 1213–1231 (2018).

60.	G. Giaever, A. M. Chu, L. Ni, C. Connelly, L. Riles, S. Véronneau, S. Dow, A. Lucau-Danila, K. Anderson, B. André, A. P. Arkin, A. Astromoff, M. El-Bakkoury, R. Bangham, R. Benito, S. Brachat, S. Campanaro, M. Curtiss, K. Davis, A. Deutschbauer, K.-D. Entian, P. Flaherty, F. Foury, D. J. Garfinkel, M. Gerstein, D. Gotte, U. Güldener, J. H. Hegemann, S. Hempel, Z. Herman, D. F. Jaramillo, D. E. Kelly, S. L. Kelly, P. Kötter, D. LaBonte, D. C. Lamb, N. Lan, H. Liang, H. Liao, L. Liu, C. Luo, M. Lussier, R. Mao, P. Menard, S. L. Ooi, J. L. Revuelta, C. J. Roberts, M. Rose, P. Ross-Macdonald, B. Scherens, G. Schimmack, B. Shafer, D. D. Shoemaker, S. Sookhai-Mahadeo, R. K. Storms, J. N. Strathern, G. Valle, M. Voet, G. Volckaert, C. Wang, T. R. Ward, J. Wilhelmy, E. A. Winzeler, Y. Yang, G. Yen, E. Youngman, K. Yu, H. Bussey, J. D. Boeke, M. Snyder, P. Philippsen, R. W. Davis, M. Johnston, Functional profiling of the Saccharomyces cerevisiae genome. *Nature*. **418**, 387–91 (2002).

61.	D. C. Zebrowski, D. B. Kaback, A simple method for isolating disomic strains of Saccharomyces cerevisiae. *Yeast*. **25**, 321–6 (2008).

62.	Y. O. Zhu, M. L. Siegal, D. W. Hall, D. A. Petrov, Precise estimates of mutation rate and spectrum in yeast. *Proc. Natl. Acad. Sci. U. S. A.* **111**, E2310-8 (2014).

63.	G. Chevereau, M. Dravecká, T. Batur, A. Guvenek, D. H. Ayhan, E. Toprak, T. Bollenbach, Quantifying the Determinants of Evolutionary Dynamics Leading to Drug Resistance. *PLoS Biol.* **13**, e1002299 (2015).

64.	R. Vaser, S. Adusumalli, S. N. Leng, M. Sikic, P. C. Ng, SIFT missense predictions for genomes. *Nat. Protoc.* **11**, 1–9 (2016).

65.	H.-H. Chou, H.-C. Chiu, N. F. Delaney, D. Segrè, C. J. Marx, Diminishing returns epistasis among beneficial mutations decelerates adaptation. *Science*. **332**, 1190–2 (2011).

66.	A. I. Khan, D. M. Dinh, D. Schneider, R. E. Lenski, T. F. Cooper, Negative epistasis between beneficial mutations in an evolving bacterial population. *Science*. **332**, 1193–6 (2011).

67.	P. Hawes, C. L. Netherton, M. Mueller, T. Wileman, P. Monaghan, Rapid freeze-substitution preserves membranes in high-pressure frozen tissue culture cells. *J. Microsc.* **226**, 182–9 (2007).

68.	E. S. Reynolds, The use of lead citrate at high pH as an electron-opaque stain in electron microscopy. *J. Cell Biol.* **17**, 208–12 (1963).

69.	J. R. Kremer, D. N. Mastronarde, J. R. McIntosh, Computer visualization of three-dimensional image data using IMOD. *J. Struct. Biol.* **116**, 71–6 (1996).

70.	H. G. Crabtree, Observations on the carbohydrate metabolism of tumours. *Biochem. J.* **23**, 536–45 (1929).

71.	H. Sies, D. P. Jones, Reactive oxygen species (ROS) as pleiotropic physiological signalling agents. *Nat. Rev. Mol. Cell Biol.* **21**, 363–383 (2020).

72.	K. Bodvard, K. Peeters, F. Roger, N. Romanov, A. Igbaria, N. Welkenhuysen, G. Palais, W. Reiter, M. B. Toledano, M. Käll, M. Molin, Light-sensing via hydrogen peroxide and a peroxiredoxin. *Nat. Commun.* **8**, 14791 (2017).

73.	S. Okazaki, T. Tachibana, A. Naganuma, N. Mano, S. Kuge, Multistep Disulfide Bond Formation in Yap1 Is Required for Sensing and Transduction of H2O2 Stress Signal. *Mol. Cell*. **27**, 675–688 (2007).

74.	A. Delaunay, D. Pflieger, M.-B. Barrault, J. Vinh, M. B. Toledano, A Thiol Peroxidase Is an H2O2 Receptor and Redox-Transducer in Gene Activation. *Cell*. **111**, 471–481 (2002).

75.	A. Hori, M. Yoshida, T. Shibata, F. Ling, Reactive oxygen species regulate DNA copy number in isolated yeast mitochondria by triggering recombination-mediated replication. *Nucleic Acids Res.* **37**, 749–761 (2009).

76.	C. M. Grant, F. H. MacIver, I. W. Dawes, Mitochondrial function is required for resistance to oxidative stress in the yeast Saccharomyces cerevisiae. *FEBS Lett.* **410**, 219–222 (1997).

77.	A. Couce, O. A. Tenaillon, The rule of declining adaptability in microbial evolution experiments. *Front. Genet.* **6** (2015), doi:10.3389/fgene.2015.00099.
